# Small grassland patches are hotspots for medicinal plants and associated phytochemical diversity in European agricultural landscapes

**DOI:** 10.1101/2025.02.28.640782

**Authors:** Rita Engel, Orsolya Valkó, Attila Lengyel, Ádám Bede, Balázs Deák

## Abstract

- Medicinal plants are threatened by overcollection and habitat loss due to human activities. In this study, we aimed to reveal how ancient earthen burial mounds covered with successional grasslands of different ages can contribute to conservation of medicinal plants and related phytochemical diversity in European agricultural landscapes.
- Using vegetation data collected from 166 mounds in Hungary, we identified the medicinal plant species present at these sites, and based on multiple literature sources, provided the secondary metabolite profiles of the medicinal plants and plant communities. By using generalized linear mixed models and redundancy analysis we investigated how the age of the vegetation, percentage of croplands around the sites, site area, and soil properties influence the diversity of medicinal plants and associated phytochemicals.
- We found that mounds provide safe havens for a wide range of medicinal plants and secondary metabolites. Our results showed that grassland-related medicinal plants producing iridoids and diterpenes were typical on mounds with older grasslands. A large proportion of croplands in the neighboring landscape and high soil pH supported generalist medicinal plants producing nitrogen-containing and sesquiterpene related secondary metabolites. Higher soil nitrogen, phosphorus, and potassium content favored medicinal plants producing lignans, glycosides, and organic acids.
- Our results suggest that the high phytochemical diversity is supported by the occurrence of various compounds facilitating the adaptation of medicinal plants to the various environmental stressors and biotic factors, such as competition, herbivory and pathogens, present in sites embedded in agricultural landscapes.

**Societal Impact Statement:** Besides being important components of landscape-level biodiversity, medicinal plants are essential resources for traditional and modern healthcare. However, in parallel with biodiversity loss driven by human activities, populations of medicinal plants have also declined. By maintaining connections between nature, culture, and people, sacred natural sites can help counteract this trend. We studied the potential of ancient burial mounds to maintain populations of medicinal plants and related phytochemical diversity in European agricultural landscapes. Our findings highlight the importance of conserving grassland vegetation and associated biodiversity at these sites, which is crucial for preserving a key ecosystem service in intensively managed landscapes.

## 1 INTRODUCTION

In many parts of the world, especially in developing countries, healthcare is still largely based on traditional remedies and, in particular, on the use of medicinal plants (World Health Organization, 2013). Approximately 60% of the global population uses traditional medicines, and in certain countries, these medicines are integral pillars of the public health system and are also used in modern healthcare and in the pharmaceutical industry (Romanelli et al., 2015; World Health Organization, 2013). Besides their direct effect on human health, as an integral parts of nature, medicinal plants also contribute to human well-being indirectly by supporting landscape aesthetics and related outdoor recreation (nature-based tourism) (McMichael et al., 2005). Furthermore, medicinal plants provide important food sources for pollinators directly benefiting their populations (Gibson & Colla, 2023; McMichael et al., 2005).

The biodiversity of plants, and specifically of medicinal plants, is being lost around the world at an unprecedented rate, largely as a result of human activities (Dudley et al., 2009; Theodoridis et al., 2024). In human-transformed landscapes, plant biodiversity is often restricted to a few protected areas; however, in many unprotected landscapes, there are alternative ways of protection by connecting cultural landscape values with biodiversity conservation (Deák et al., 2023; Dudley et al., 2009). In many cases, religious, spiritual, and cultural functions connected to a certain site can effectively support the preservation of natural and semi-natural habitats, and the species confined to them (Zannini et al., 2021). These sites, known as sacred natural sites (SNSs), represent areas of spiritual significance can exist in various forms of natural (e.g., sacred groves, mountains, caves, and rivers) and anthropogenic (churchyards, old cemeteries, ancient fortifications or burial sites) features (Zannini et al., 2021). Due to taboos and extensive management associated with SNSs, they could avoid land transformation actions and thus they are often characterized by high biodiversity and act as refuge for rare and endemic species (Deák et al., 2023). Apart from conserving biodiversity in general, these sites also have a potential role in the conservation of medicinal plants (Zannini et al., 2021). In the scientific literature, we can find examples for the positive effects of SNSs on medicinal plant conservation in Asia and Africa, such as sacred groves in Kodagu, South India (Boraiah et al., 2003), sacred forests in North Thailand (Junsongduang et al., 2013), and in Sidama, Ethiopia (Doffana, 2017). Despite the potential role of SNSs in maintaining populations of medicinal plants, it is a rarely studied topic in the developed countries with temperate climates. However, their potential importance is shown by a study about ancient Lithuanian hillforts (protected archaeological sites) that these sites serve as genetic reserves for medicinal plants, supporting their in-situ conservation (Labokas & Karpavičienė, 2023).

In the continental steppes of Eurasia, with an estimated half a million documented occurrences, ancient burial mounds (so-called “kurgans”) are probably the most widespread ancient, manmade SNSs (Apostolova et al., 2022; Deák et al., 2016; Moysiyenko et al., 2014). These hill-like earthen structures, with heights generally ranging from half a meter to five meters, were built during the Copper, Bronze, and Iron Ages by steppe nomad cultures such as the Yamnayas, Scythians, Sarmatians or later ancient Turkic nomads (Deák et al., 2023). Although originally built for burial purposes, over millennia subsequent generations have used kurgans for secondary burials even till present days, they have served as grounds for religious objects and rituals, and often have played important role in folklore (Deák et al., 2023). In many cases, special bound of local populations prevented detrimental land use changes, such as ploughing or infrastructure developments (Deák et al., 2024). Mounds still covered with semi-natural grasslands function as safe havens and biodiversity hotspots for grassland-related taxa, providing an additional pillar for nature conservation in human transformed landscapes (Deák et al., 2023). This is especially important, as steppes and associated species become seriously threatened worldwide mostly due to the expansion and intensification of agricultural lands (Wesche et al., 2016). Despite their cultural and conservation importance, during the past centuries, many kurgans have been ploughed and converted to cropland (Deák et al., 2020; Moysiyenko et al., 2014). Recognizing the importance of kurgans in the preservation of the traditional landscape and related biodiversity, in Hungary, kurgans are protected by law (Nature Conservation Act 1996/LIII), and farmers applying for agri-environmental schemes associated with the Common Agricultural Policy of the EU must avoid any activities related to arable farming on them (Deák et al., 2023). As a result, the spontaneous regeneration of grassland vegetation has begun on many formerly ploughed kurgans, leading to the co-existence of kurgans covered with millennia-old grasslands and successional vegetation, both potentially holding a high diversity of medicinal plants.

The medical effect of medicinal plants is related to their secondary metabolite content (Buchanan et al., 2015). The variety of secondary metabolites produced by medicinal plants forms the basis of the landscape-level phytochemical diversity (Defossez et al., 2021). Secondary metabolites are organic compounds that are not directly involved in plant growth and development. To date, more than 200,000 secondary metabolites have been identified, and these can be classified according to their biosynthetic origin and chemical structure (Buchanan et al., 2015). These compounds play an important role in supporting plants in their establishment and survival under specific environmental conditions (protection against UV radiation, improvement of stress tolerance against drought, heat, freezing, salt) (Kessler & Kalske, 2018; Pant et al., 2021). They also have specific functions in plant-animal interactions such as pollinator attraction and seed dispersal (Kessler & Kalske, 2018; Pant et al., 2021).

Through similar biological functions, plant secondary metabolites can influence landscape-scale ecosystem processes (Wetzel & Whitehead, 2020) and play fundamental roles in vegetation dynamics by influencing community-level biotic interactions through direct chemical defense (antifeedants, allelochemicals) or chemical signaling (mediating interactions with antagonists such as herbivores and pathogens, and mutualists such as mycorrhizal fungi) (Engel et al., 2016; Kessler & Kalske, 2018). As an example, medicinal plants was shown to gain advantages during early succession through the production of secondary metabolites (Kessler & Kalske, 2018). For instance, tannins produced by balsam poplar (*Alnus incana* ssp. *tenuifolia*) inhibits nitrogen-fixing bacteria and nitrogen mineralization in soils, creating environmental conditions (tannins released by balsam poplar inhibited nitrogen-fixing bacteria) that support a successional shift from grey alder (*Alnus incana* ssp. *tenuifolia*) to poplar dominance (Schimel et al., 1998).

The synthesis of secondary metabolites is strongly influenced by climatic and topographic conditions (Pant et al., 2021; Tiwari et al., 2023). For example, in a study the positive effects of slope, altitude, total nitrogen and organic carbon content of the soil on the hypericin content of *Hypericum perforatum* were reported (Saffariha et al., 2021). The strong relationship between environmental conditions and secondary metabolite production may influence the habitat preferences and geographic distribution of medicinal plants producing different types of compounds. However, few studies have investigated this topic (Hazlett & Sawyer, 1998; Tiwari et al., 2023). In a related study, it was found, that alkaloid-rich species were more abundant in water-enriched habitats in Colorado (Hazlett & Sawyer, 1998).

Using data collected in our regional-scale survey of 166 sites, we aimed to reveal how ancient burial mounds, a typical example of SNSs of the continental steppe region, can contribute to the preservation of medicinal plants and related phytochemical diversity in a European agricultural landscape. We also aimed to investigate the factors that influence these diversity patterns. Our study questions were as follows: (i) Do kurgans holding successional and millennia-old grasslands have the potential to play role in the maintenance of medicinal plant populations in transformed agricultural landscapes? (ii) Do the age of the vegetation, land use intensity (expressed by the percentage of arable land around the kurgans), the area of the kurgans, and soil properties impact the species richness and abundance of medicinal plants and the related phytochemical diversity?

## 2 METHODS

### 2.1 Study area and sites

Our study area is located in the lowlands of the Hungarian Great Plain (East Hungary) and covers approximately 21,000 km^2^. The climate of the area is continental (mean annual precipitation: 545 mm; mean annual temperature: 10.2 °C). In historical times, the dominant vegetation of the area was forest steppe, composed of a complex mosaic of small patches of woody vegetation embedded in a matrix of dry to wet grasslands. The most species-rich habitat of this habitat complex is the loess steppe, which holds a high biodiversity of grassland specialist grass, sedge, and forb species, including many medicinal plants (Deák et al., 2021). For today, due to the expansion of arable lands and infrastructure associated with residential and industrial areas, a considerable proportion of the natural vegetation has been lost. Loess steppes have been the most severely affected, as their fertile chernozemic soil is suitable for crop production, making them primary targets of land transformation throughout history. Today, only a few percent of their original stands remain intact (Biró et al., 2018). In the agricultural landscapes of the study area, these stands are mostly found on road verges and on kurgans (Deák et al., 2021).

For this study, we selected 166 isolated kurgans that were embedded in agricultural areas and covered with either successional grasslands or ancient undisturbed grasslands. The mean area of the mounds was 3,309 (± 2,345 SD) m^2^, and their height was 2.9 (± 1.5 SD) meters. The successional grasslands represented a large age gradient, ranging from 1 to 140 years after the cessation of ploughing. We also considered kurgans that have been continuously covered with ancient, undisturbed grasslands since the Bronze Age (approximately for 5000 years). To estimate the age of successional grasslands, we used cartographic data from the Second (1858-1864) and Third (1881-1884) Military Survey of the Hungarian Empire, the Military Survey of the Hungarian People’s Republic (1956-1975), and the current topographical map of Hungary (Institute and Museum of Military History, Budapest), archived aerial photographs (source: www.fentrol.hu), and satellite data (time series of satellite images from Google Earth; Google Earth 2024). As kurgans serve as good orientation points in the flat landscapes, they and their landcover have generally been included on topographical maps for centuries, which enables accurate estimation of their successional age. Younger successional grasslands on mounds were mostly dominated by weedy fallow vegetation, which, over time, shifted towards a vegetation composed of species typical of loess grassland vegetation. The non-ploughed, millennia-old kurgans were covered with diverse grassland vegetation.

### 2.2 Vegetation sampling

The vegetation sampling spanned two years (2021–2022), with each site surveyed once during the peak vegetation period (May or June). At each site, we documented the vascular plant species present and visually estimated the percentage cover of each species. To compile the species lists, we performed a comprehensive visual survey of the entire site. Following a standardized protocol, three surveyors dedicated 30 minutes per 0.1 ha during each survey. Plant names nomenclature follows Király et al. (2009).

### 2.3 Habitat mapping

Simultaneously with the botanical survey, we mapped arable lands (as a proxy for the level of landscape transformation) within a 300-meter buffer surrounding each site. This was achieved using satellite imagery from Google Maps (2024) and Bing Maps (2024). We digitized the maps prepared in the field and calculated the area of arable lands using QGIS software (QGIS Development Team, 2023).

### 2.4 Soil sampling

At the time of the botanical survey, we collected soil from the top 10 cm at five random points on each mound, resulting in a 1,000 cm^3^ composite soil sample. The samples were air-dried and homogenized, and the phosphorus (P), nitrogen (N), and potassium (K) content and pH of the soil were analysed in an accredited laboratory (reference number: NAH-1-1437/2018).

### 2.5 Data processing

#### 2.5.1 Plant functional groups

Based on (Király et al., 2009), we classified species into perennial and short-lived categories. We also assigned them based on their habitat affinity to grassland-related and generalist groups. A species was assigned to the grassland-related group, if it occurred mainly in grassland habitats in the study area (Király et al., 2009). All other species were assigned to the generalist group, which contained species typical of semi-natural habitats but without a definite habitat preference and species with a weed strategy.

#### 2.5.2 Medicinal plants and secondary metabolites

A plant species was considered a medicinal plant if (i) it was included in a pharmacopoeia (e.g., Hungarian Pharmacopoeia, European Pharmacopoeia), (ii) it was listed in a herbal monograph, or (iii) it was mentioned as a medicinal plant in a relevant peer-reviewed publication (e.g., Journal of Ethnopharmacology, Planta Medica) confirming its medicinal properties, its traditional medicinal use, or the presence of secondary metabolites responsible for its medicinal effects. Table S1 lists the medicinal plants found on the studied kurgans, their secondary metabolites, their medicinal effects, and the relevant publications mentioning the species.

Based on the literature, secondary metabolites produced by medicinal plants were collected and classified into 33 groups: (1) flavonoids, (2) phenolic compounds, (3) tannins, (4) essential oils, (5) alkaloids, (6) plant steroids, (7) triterpenes, (8) polysaccharides, (9) vitamins, (10) other non-alkaloid nitrogen-containing compounds, (11) saponins, (12) coumarins, (13) organic acids, (14) mineral nutrients, (15) fatty acids, (16) plant proteins, (17) iridoids, (18) lignans, (19) sesquiterpenes, (20) sesquiterpene lactones, (21) carotenoids, (22) resins, (23) diterpenes, (24) quinones, (25) steroidal glycosides, (26) glucosinolates, (27) xanthones, (28) naphthalene and derivatives, (29) triglycerides, (30) phloroglucinol derivatives, (31) cyanogenic glucosides, (32) monosaccharides, and (33) phyto-cannabinoids. We calculated the Simpson’s Index of Diversity of these secondary metabolite groups on each mound taking into account the abundance of species containing the individual metabolite groups. Diversity scores, as well as the abundance-weighted frequency of the secondary metabolite groups, were used as response variables in the statistical analyses. Additional details and references regarding the classification of secondary metabolites are provided in Table S2.

We conducted an extensive literature search but did not carry out laboratory analyses to characterize the secondary metabolites of the plants. Thus, we could only estimate the potential presence of the secondary metabolites in the medicinal plants found on the kurgans. It should be noted that, due to the specific habitat conditions, the metabolite production of a certain species may differ from what is described in the literature. However, we believe that the use of literature data provides a good estimation of the phytochemical diversity patterns on the studied kurgans.

#### 2.5.3 Statistical analysis

We used generalized linear mixed models (GLMM) to reveal relationships between community-level descriptors of diversity and composition as response variables, and landscape and soil variables as explanatory variables. More specifically, our response variables were the richness and cover of medicinal plants, the richness and cover of short-lived, perennial, grassland-related and generalist medicinal plants, and Simpson diversity of secondary metabolites. The explanatory variables included the grassland ages, the percentage of cultivated land in the 300-m buffer, the area of the kurgans, soil pH, and soil nitrogen, phosphorous and potassium content. We applied a logarithmic transformation on grassland age in order to reduce the skewness caused by large values and to make the data distribution more symmetrical. We tested all predictors for multicollinearity using variance inflation factors (VIF). Since the VIF was lower than 1.5 in each case (multicollinearity being negligible), we considered all explanatory variables for the statistical analyses. First, all explanatory variables were included in a maximal model. Then, the model was optimized using stepwise backward variable selection based on AIC values. We tested the effects of the variables in the final model by Wald tests. For fitting models on species richness values, we used a negative binomial distribution to account for overdispersion in the data. Percentage cover scores of species groups were transformed to have a range from 0.001 to 0.999, and then we used the beta distribution family for them in the models. For Simpson’s index of diversity scores calculated for the abundance-weighted frequency of secondary metabolites, we fitted models with a normal distribution.

#### 2.5.4 Redundancy analysis (RDA)

We assessed and visualized the effect of the predictors used in the GLMMs on the abundance weighted frequency of the 33 groups of secondary metabolites (dependent variables) using redundancy analysis (RDA) and stepwise backward selection of the predictors. To calculate the abundance-weighted frequency of secondary metabolites, we used the percentage cover scores of the plant species as weights. We performed Hellinger transformation on the frequency scores, which assigns low weights to rare metabolites and allows the use of RDA even in the case of a long gradient in the data. We assessed the significance of model terms by a permutation test using 999 random permutations.

All calculations were carried out in R (R Core Team, 2021). To calculate VIFs, we used the ‘faraway’ package in R (Faraway, 2016). For GLMMs with stepwise backward selection, we used the glmmTMB (Brooks et al., 2017) and MuMIn packages (Burnham & Anderson, 2002). For calculating RDA we used the ‘vegan’ package in R (Oksanen et al., 2019).

## 3 RESULTS

### 3.1 Medicinal plants on kurgans

On the 166 kurgans involved in our study, we detected a total of 164 medicinal plant species, including 42 grassland-related species. Fifty-five percent of the species were perennial, and 45% were short-lived. The number of medicinal plants found per kurgan ranged from 5 to 34 (mean= 17.6 ± 5.8 SD, median = 18). The five most frequent medicinal plants present on the mounds were *Convolvulus arvensis*, *Capsella bursa-pastoris*, *Elymus repens*, *Cirsium arvense*, and *Achillea collina* (Figure 1). Apart from *Achillea collina*, the most frequent grassland-related medicinal plants were *Podospermum canum*, *Tragopogon dubius*, *Galium verum*, and *Epilobium tetragonum* (Figure 1).

**Figure 1.**
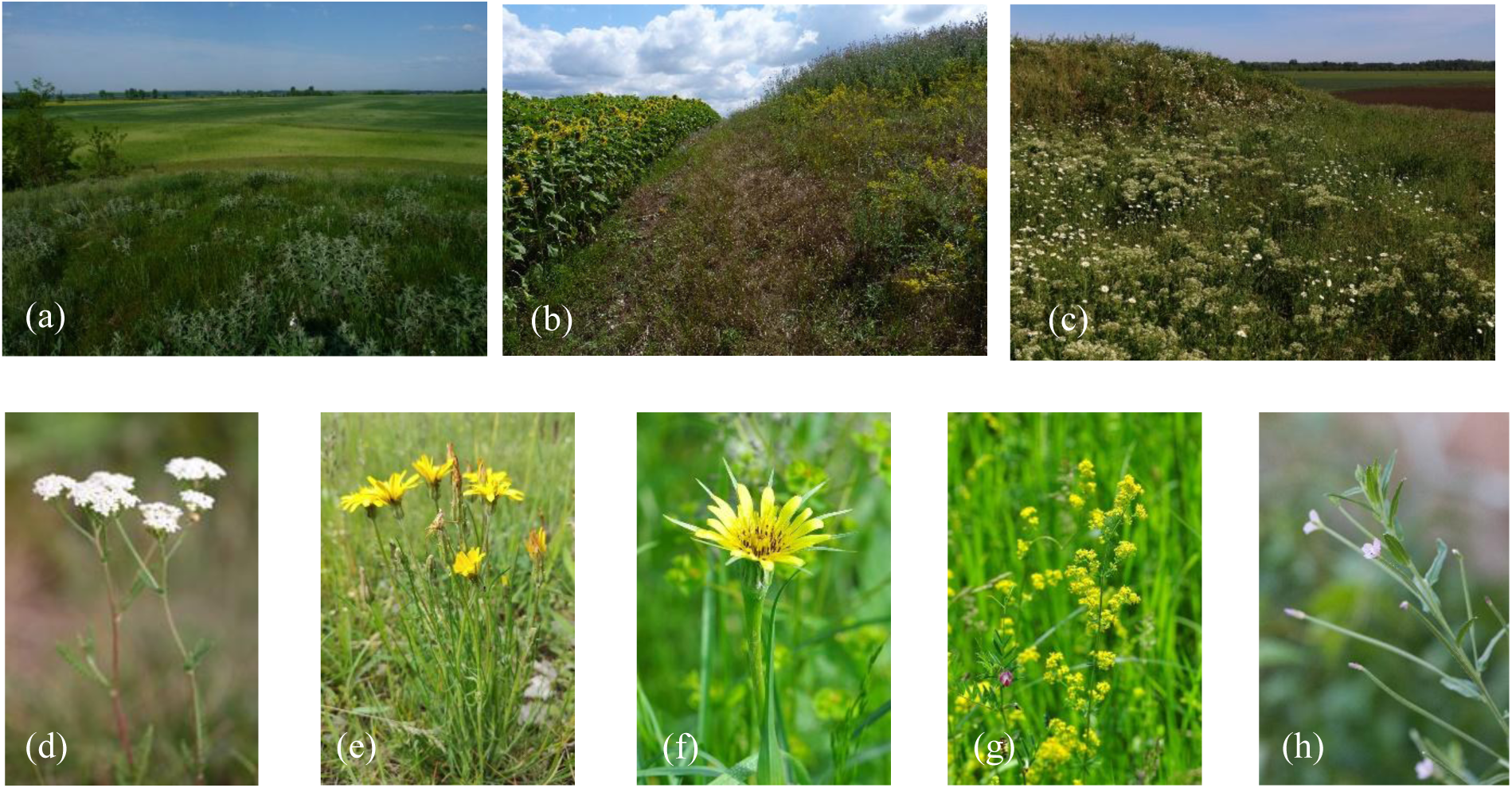
(a) Kurgan with a pristine grassland, (b) old field and (c) young fallow; The most frequent grassland-related medicinal plants on kurgans: (d) *Achillea collina* (102 mounds), (e) *Podospermum canum* (79 mounds), (f) *Tragopogon dubius* (66 mounds), (g) *Galium verum* (64 mounds), (h) *Epilobium tetragonum* (42 mounds) (photos by Balázs Deák a, b, c, d and Attila Lengyel e, f, g, h)

### 3.2 Factors affecting medicinal plants

Grassland ages were positively related to the species richness and cover of grassland-related and perennial medicinal plants (Table 1). Kurgans surrounded by a higher percentage of cultivated land were characterized by a lower cover of medicinal plants due to the lower cover of perennial medicinal plants. Grassland-related medicinal plants were represented by lower species richness on these kurgans (Table 1). Kurgans with a larger area held a greater species richness and cover of medicinal plants. This pattern was associated with the higher cover of generalist medicinal plants in larger mounds. The cover of grassland-related medicinal plants was not affected by the area of the kurgans. Kurgans with higher soil pH held higher species richness of short-lived and generalist medicinal plants. Lower soil pH was associated with higher cover of grassland-related medicinal plants. High soil nitrogen content supported higher cover of short-lived and generalist medicinal plants. Low soil nitrogen content was related to higher species richness of grassland-related medicinal plants. Soil phosphorus (P) and potassium (K) content did not affect the species richness and cover of medicinal plants.

**Table 1.**
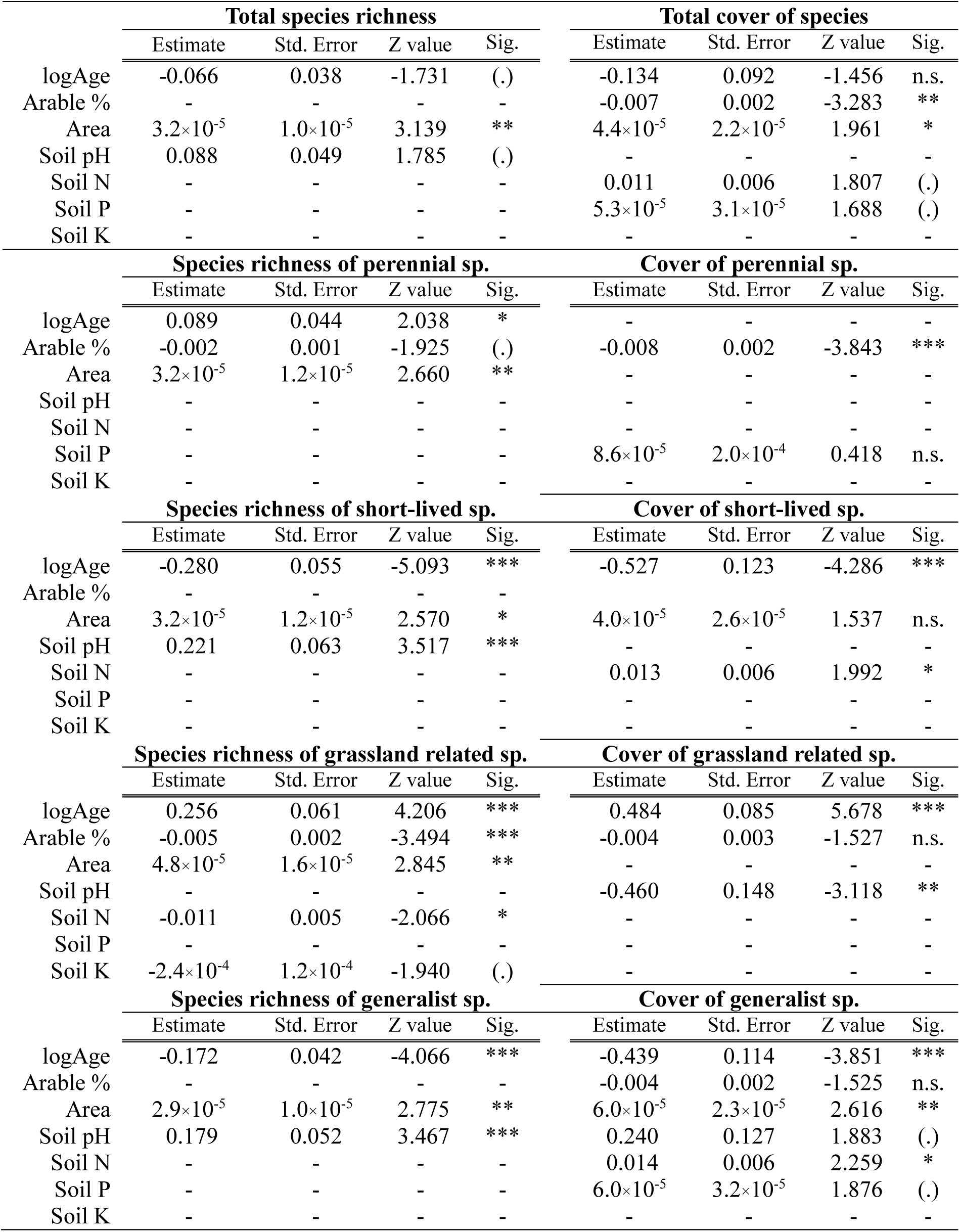
Effects of abiotic environmental factors on the species richness and cover of medicinal plants (generalized linear mixed models with backward selection). Abbreviations of the predictors: logAge –grassland age, Arable % – percentage proportion of cultivated land around the kurgans within a 300-m buffer, Area – area of the kurgan, Soil pH– soil pH value, Soil N– soil nitrogen content. Notation of significant results: * p≤ 0.05, ** p≤ 0.01, *** p≤ 0.001, (.) marginal significance, n.s. not significant.

### 3.3 Phytochemical diversity

Based on the collected literature data, medicinal plants present in the surveyed mounds contained a total of 33 secondary metabolite groups, of which 23 were represented in grassland-related medicinal plant species. Eleven secondary metabolite groups (alkaloids, coumarins, flavonoids, other non-alkaloid nitrogen-containing compounds, phenolic compounds, plant proteins, saponins, plant steroids, tannins, triterpenes, and vitamins) were represented on all mounds (Table 2). In the medicinal plants, flavonoids, other phenolic compounds, essential oils, tannins, and steroids were the dominant secondary metabolite groups (Table 2). On the kurgans, the five most frequent secondary metabolite groups were flavonoids, other phenolic compounds, tannins, essential oils, and alkaloids (Table 2).

**Table 2.** Frequency and occurrence of secondary metabolites produced by medicinal plants found on the kurgans. Pooled Frequency (%) = (total occurrence of all secondary metabolites / occurrence of the secondary metabolite) ×100; Number of species: number of medicinal plants containing the certain secondary metabolite; Occurrence number: number of mounds where the certain secondary metabolite occurred.

|  | Secondary metabolites | Pooled frequency (%) | Number of species | Occurrence number |
| --- | --- | --- | --- | --- |
| 1 | Flavonoids | 18.99 | 108 | 166 |
| 2 | Phenolic compounds | 8.38 | 69 | 166 |
| 3 | Tannins | 6.44 | 42 | 166 |
| 4 | Essential oils | 5.97 | 48 | 164 |
| 5 | Alkaloids | 5.31 | 27 | 166 |
| 6 | Plant steroids | 5.14 | 28 | 166 |
| 7 | Triterpenes | 5.12 | 27 | 166 |
| 8 | Polysaccharides | 4.07 | 21 | 155 |
| 9 | Vitamins | 3.83 | 17 | 166 |
| 10 | Non-alkaloid nitrogen cont. comp. | 3.65 | 14 | 166 |
| 11 | Saponins | 3.54 | 24 | 166 |
| 12 | Coumarins | 2.87 | 12 | 166 |
| 13 | Organic acids | 2.80 | 14 | 148 |
| 14 | Mineral nutrients | 2.50 | 15 | 142 |
| 15 | Fatty acids | 2.44 | 15 | 134 |
| 16 | Plant proteins | 2.44 | 8 | 166 |
| 17 | Iridoids | 2.39 | 19 | 137 |
| 18 | Lignans | 1.93 | 12 | 130 |
| 19 | Sesquiterpenes | 1.88 | 6 | 125 |
| 20 | Sesquiterpene lactones | 1.82 | 13 | 131 |
| 21 | Carotenoids | 1.62 | 9 | 128 |
| 22 | Resins | 1.60 | 2 | 165 |
| 23 | Diterpenes | 1.20 | 18 | 84 |
| 24 | Quinones | 0.86 | 11 | 82 |
| 25 | Steroidal glycosides | 0.81 | 5 | 88 |
| 26 | Glucosinolates | 0.74 | 3 | 83 |
| 27 | Xanthones | 0.68 | 1 | 79 |
| 28 | Naphthalene and derivatives | 0.51 | 5 | 56 |
| 29 | Triglycerides | 0.23 | 7 | 25 |
| 30 | Phloroglucinol derivatives | 0.10 | 1 | 12 |
| 31 | Cyanogenic glucosides | 0.07 | 1 | 8 |
| 32 | Monosaccharides | 0.03 | 3 | 4 |
| 33 | Phyto-cannabinoids | 0.03 | 1 | 4 |

The phytochemical diversity was significantly higher on kurgans with higher soil potassium content. Conversely, high soil nitrogen content resulted in a lower phytochemical diversity (Table 3).

**Table 3.** Effects of abiotic environmental factors on the phytochemical diversity of medicinal plants based on Simpson diversity index (Generalized Linear Mixed Models). Abbreviations: logAge – grassland age, Arable % – percentage proportion of cultivated land around the kurgans within a 300-m buffer, Soil pH– soil pH value, Soil N– soil nitrogen content, Soil P– soil phosphorus content, Soil K– soil potassium content. Notation of significant results: * p≤ 0.05, ** p≤ 0.01, *** p≤ 0.001, (.) marginal significance.

|  | Simpson diversity Index |  |  |  |
| --- | --- | --- | --- | --- |
|  | Estimate | Std. Error | Z value | Sig. |
| logAge | - | - | - | - |
| Arable % | $1.6 \times 10^{-4}$ | $9.9 \times 10^{-5}$ | 1.706 | (.) |
| Area | - | - | - | - |
| Soil pH | - | - | - | - |
| Soil N | $-9.6 \times 10^{-4}$ | $2.9 \times 10^{-4}$ | -3.230 | ** |
| Soil P | $-3.3 \times 10^{-6}$ | $1.9 \times 10^{-6}$ | -1.762 | (.) |
| Soil K | $2.5 \times 10^{-5}$ | $8.9 \times 10^{-6}$ | 2.832 | ** |

The results of the redundancy analysis (RDA) showed that the grassland age, the soil nitrogen, phosphorus and potassium content had a significant effect on the abundance of different secondary metabolites produced by medicinal plants. (Table 4).

**Table 4.** Effects of abiotic environmental factors on the abundancy of secondary metabolites (redundancy analysis using backward selection). Abbreviations: logAge –grassland age, Arable % – percentage proportion of cultivated land around the kurgans within a 300-m buffer, Soil pH– soil pH value, Soil N– soil nitrogen content, Soil P– soil phosphorus content, Soil K– soil potassium content. Notation of significant results: * p≤ 0.05, ** p≤ 0.01, *** p≤ 0.001, (.) marginal significance.

| Abundance of secondary metabolites |  |  |  |  |
| --- | --- | --- | --- | --- |
|  | Df. | Variance | F | Sig. |
| logAge | 1 | 0.011 | 11.060 | *** |
| Arable % | 1 | 0.002 | 1.826 | (.) |
| Soil pH | 1 | 0.002 | 2.223 | * |
| Soil N | 1 | 0.005 | 4.681 | *** |
| Soil P | 1 | 0.003 | 2.719 | * |
| Soil K | 1 | 0.004 | 4.083 | *** |

The occurrence of diterpenes (23), iridoids (17), carotenoids (21), and essential oils (4) were typical on kurgans covered with older successional grasslands. Saponins (11), coumarins (12), steroids (6), alkaloids (5), proteins (16), other non-alkaloid nitrogen-containing compounds (10), and resins (22) were typical on kurgans surrounded by a larger proportion of arable lands. The area of the mounds had no significant impact on secondary metabolites. Kurgans with alkaline soils held a higher abundance of sesquiterpenes (19), sesquiterpene lactones (20), and fatty acids (15). Higher soil nitrogen levels supported a higher abundance of lignans (18) and essential oils (4). We found that on kurgans with higher soil phosphorus content, polysaccharides (8), minerals (14), glucosinolates (26), flavonoids (1), fatty acids (15), and steroidal glycosides (25) were typical. Higher soil potassium level resulted in a greater abundance of tannins (3), triterpenes (7), phenolic compounds (2), and organic acids (13) (Figure 2).

**Figure 2.**
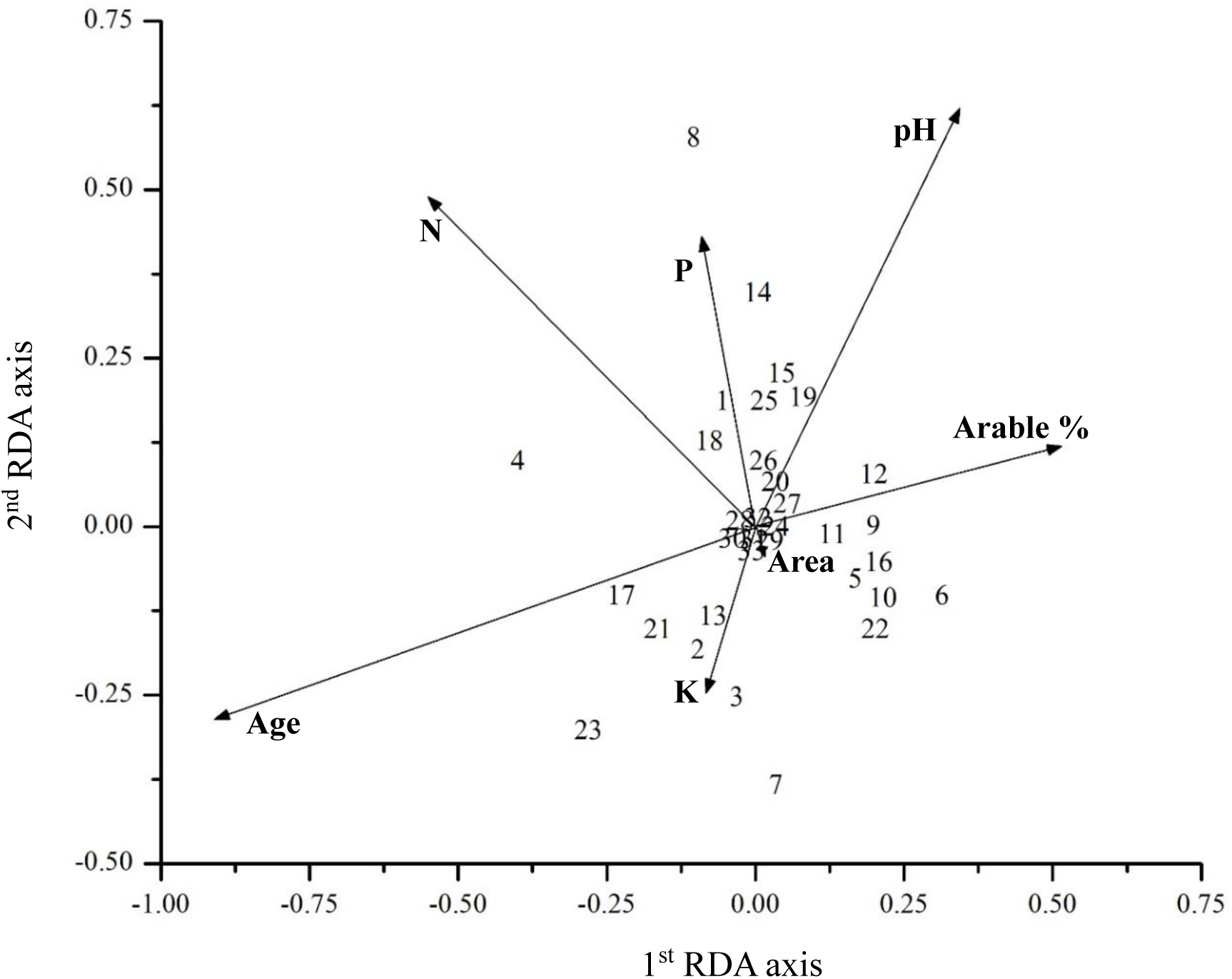
The effect of grassland age (Age), percentage of cultivated land around the mounds (Arable %), soil nitrogen (N), phosphorus (P) and potassium (K) content, and pH on the phytochemical composition of plant assemblies (redundancy analysis with stepwise backward variable selection). Notation of secondary metabolite groups: 1 – flavonoids, 2 – phenolic compounds, 3 – tannins, 4 – essential oils, 5 – alkaloids, 6 – plant steroids, 7 – triterpenes, 8 – polysaccharides, 9 – vitamins, 10 – other non–alkaloid nitrogen-containing compounds, 11 – saponins, 12 – coumarins, 13 – organic acids, 14 – mineral nutrients, 15 – fatty acids, 16 – plant proteins, 17 – iridoids, 18 – lignans, 19 – sesquiterpenes, 20 – sesquiterpene lactones, 21 – carotenoids, 22 – resins, 23 – diterpenes, 24 – quinones, 25 – steroidal glycosides, 26 – glucosinolates, 27 – xanthones, 28 – naphthalene and derivatives, 29 – triglycerides, 30 – phloroglucinol derivatives, 31 – cyanogenic glucosides, 32 – monosaccharides, 33 – phyto– cannabinoids. Variance explained by the 1^st^ and 2^nd^ RDA axes were 68.7% and 17.6% respectively.

## 4 DISCUSSION

We found that kurgans situated in agricultural landscapes hold a high diversity of medicinal plants. This diversity is supported by the fact that kurgans covered with successional grasslands represent a high variability both in environmental factors and the composition of plant communities. In connection with the high diversity of medicinal plants, kurgans also act as phytochemical hotspots in the landscape.

### 4.1 Grassland age

We found that the older the grassland on the mounds, the higher the abundance of secondary metabolites (such as iridoids and diterpenes) that enhance the heat and drought tolerance of medicinal plants. A possible reason for this is that during succession, the green biomass considerably decreases over time, which reduces the level of self-shading in the vegetation (Van Loon et al., 2014). In older grasslands, grassland-related medicinal plants gain dominance, and these species, adapted to open habitat conditions, are more tolerant to heat and drought, partly due to the secondary metabolites produced by them (Pant et al., 2021). In relation to this phenomenon, the study by Wang et al. (2010) showed that the drought stress increased the accumulation of iridoid glycosides in the roots of *Scrophularia ningpoensis*, which provides protective activities against oxidative stress induced by water deficit conditions (Wang et al., 2010). In line with these findings, the higher presence of iridoids (17) on kurgans with older vegetation can be closely related to the presence of *Galium verum*, a typical perennial grassland-related herb species in the region. Among the 19 iridoid-containing species, the frequency and cover of the perennial grassland-related *Galium verum* were remarkably high compared to the other iridoid-producing medicinal plants. *Galium verum* may produce iridoids because of their antioxidant (free radical scavenging) activity (Cuendet et al., 1997), which might support the plant in adapting arid, highly UV-exposed habitats such as the dry grasslands of the kurgans.

The greater occurrence of diterpenes (23) in older grasslands is likely due to the higher species richness and cover of perennial grassland-related medicinal plants, as most (87.5%) of plants producing diterpenes are perennial. In our studied sites the most abundant diterpene-containing species were from the *Salvia* genus. A study by Munné-Bosch et al. (2001) identified the role of carnosic acid, a diterpene molecule, in protecting the leaves of *Salvia officinalis* from drought-induced oxidative stress. They found that carnosic acid levels in the leaves decreased as drought progressed, and at the same time, an increase in its oxidation products was detected. This antioxidant protective mechanism of diterpenes is responsible for preventing drought-induced damage in plants (Munné-Bosch et al., 2001). In this way, the production of diterpenes can also be part of the adaptation strategy to arid habitats.

### 4.2 Percentage of arable land

Medicinal plants present on the kurgans surrounded by high percentage of arable land produce a wide variety of nitrogen-containing compounds, such as alkaloids (5), proteins (16), other non-alkaloid nitrogen-containing-compounds (10), saponins (11), coumarins (12), plant steroids (6), vitamins (9), and resins (22). These compounds play an important role in the defense system of the plants. Most of them (alkaloids, coumarins, plant steroids, saponins) act as antifeedant chemicals due to their toxicity (Purrington, 2003). These secondary metabolites contribute to the increased tolerance and adaptation of medicinal plants with a weed strategy, which is supported by the finding that *Convolvulus arvensis*, the most frequent generalist medicinal plant on the mounds, which generally have a high cover produces all of these compounds.

### 4.3 Area of kurgans

Kurgans can act as hotspots of phytochemical diversity in the agricultural landscape, regardless of their size, as their area did not affect the diversity of secondary metabolites provided by the medicinal plants present. However, larger mounds had a richer pool of medicinal plants, due to the larger area, which can support the coexistence of more medicinal plants with a higher cover. Nevertheless, the uniformly high phytochemical diversity of the mounds, regardless of their size, may be due to differences in the species composition of the vegetation.

### 4.4 Soil pH value

We found that the soil pH positively affected the abundance of sesquiterpene-related compounds. On kurgans with higher soil pH, medicinal plants producing sesquiterpenes (19, e.g., *Anthemis arvensis*, *Onopordum acanthium*, *Taraxacum officinale*) and sesquiterpene lactones (20, e.g., *Crepis biennis*, *Descurainia sophia*, *Stenactis annua*) were typical. Most of these medicinal plants (89%) were generalist species. Sesquiterpene lactones are bitter compounds forming a large group of secondary metabolites. They are present in several plant families but they are predominant in the Asteraceae family (Padilla-Gonzalez et al., 2016). We observed the same pattern, with 93% of plants producing sesquiterpene lactones on the studied kurgans belonged to the *Asteraceae* family. Sesquiterpene lactones have several ecological functions. They play a protective role against herbivorous insects and pathogens. Due to their phytotoxicity (growth regulatory activities), they inhibit the growth of competing plants (Padilla-Gonzalez et al., 2016). They also act as signaling compounds in the rhizosphere to help establish (e.g. the arbuscular mycorrhizal) symbioses (Frey et al., 2024). Due to their wide range of ecological functions, the production of sesquiterpene lactones can be influenced by various environmental factors (e.g. light intensity, temperature, soil composition etc.) (Frey et al., 2024). A possible explanation for our findings may be a complex, indirect effect of more environmental factors. For example, one theory can be that, the secretion of sesquiterpene lactones facilitates the establishment of arbuscular mycorrhizal symbiosis, giving the plant better access to nutrients that are limited in high pH soils. This, in turn, enables the plant to produce more sesquiterpene lactones, thereby strengthening its defense against environmental stresses such as herbivory. Similar to our findings, a positive relationship between sesquiterpene lactone content and soil pH was found in a study investigating *Arnica montana* (*Asteraceae*) populations in grasslands of the Apuseni Mountains in Romania. They found that higher soil pH increased the number of individuals, and also the sesquiterpene lactone content of *Arnica montana* (Greinwald et al., 2022).

### 4.5 Soil nitrogen

Our results suggest that mounds with higher soil nitrogen level favored generalist medicinal plants. Nutrient rich environments in successional grasslands enhance plant growth and biomass production, leading to high levels of competition (Deák et al., 2020). In such environments, medicinal plants can gain a competitive advantage through the production of secondary metabolites such as lignans (18) and essential oils (4). Lignans can support plants by enhancing their protection against disease, herbivory, and UV radiation, and they also have a strong allelopathic effect (Ražná et al., 2021). In the case of many medicinal plants (such as *Arctium lappa, Onopordum acanthium*, and *Urtica dioica*) present on kurgans, previous studies revealed this allelopathic effect and showed their inhibitory effect on seed germination and seedling growth (Bojović et al., 2018; Suzuki et al., 2019; Watanabe et al., 2014).

Lignans may contribute to this phytotoxic effect within the complex bioactive productivity of these plants. This explains the lower phytochemical diversity associated with higher soil nitrogen levels. The allelopathic effects of the plants confined to higher soil nitrogen can cause a decrease in species richness, which is accompanied by a decrease in phytochemical diversity. Our results suggest that a high proportion of lignan producing plants on kurgans characterized by high soil nitrogen content can contribute to a general decrease in the species richness and cover of other medicinal plants by the allelopathic effect due to enhanced lignan production.

More than 70% of the essential oil producing medicinal plants found on the kurgans were generalist species, such as *Arctium lappa* (Suzuki et al., 2019). Essential oils, besides their role in pollinator attraction, anti-pathogenic, and anti-herbivorous properties, also act as allelopathic chemicals, similar to lignans (Buchanan et al., 2015).

### 4.6 Soil phosphorus

Mounds characterized by high soil phosphorus content harbored medicinal plants that produce sugar-containing secondary metabolites (e.g., glucosinolates and steroidal glycosides: *Descurainia sophia*, flavonoids: *Capsella bursa-pastoris*, *Convolvulus arvensis*, polysaccharides: *Elymus repens*), because higher phosphorus availability can cover the extra energy cost of synthesizing these secondary metabolites required for plant development in larger amounts (Neilson et al., 2013).

Glucosinolates (26) support the plant defense system against herbivores, but this protection is costly. Bekaert et al. (2012) found that glucosinolate production in *Arabidopsis thaliana* increased the photosynthetic requirements by at least 15%. Züst et al. (2011) observed a negative correlation between glucosinolate content and plant growth rates during early development of *Arabidopsis thaliana*. This confirms that defense by glucosinolates is costly for the plant because their production restrains plant growth. However, some medicinal plants can synthesize glucosinolates without any negative effect on their growth if there is a sufficient amount of available phosphorus in the soil, which is essential for the process of converting inorganic carbon into carbohydrates (sugars) during photosynthesis. The same applies to other sugar-containing secondary metabolites, such as steroidal glycosides, flavonoids, and polysaccharides, produced by medicinal plants (Neilson et al., 2013).

Steroidal glycosides (25) are toxic compounds that act mainly as feeding deterrents to protect the plants from herbivores (Purrington, 2003). This anti-herbivore effect is an important ecological strategy for medicinal plants on kurgans, as in the agricultural landscapes they are highly exposed to herbivores (mostly arthropods) from surrounding arable fields.

In nature, flavonoids (1) usually occur in a glycosylated form (flavonoid glycosides) and are widely distributed in higher plants (Karakaya, 2004). In our study, 66% of the medicinal plants found on the kurgans can potentially produce flavonoids. Flavonoids play multiple roles in plant responses to various abiotic and biotic environmental stresses. Among these, photoprotection plays a prominent role on the kurgans, as the medicinal plants growing there are exposed to strong solar radiation due to the lack of shading woody species. Flavonoids are able to reduce light-induced oxidative damage due to their UV screening properties and antioxidant functions. Furthermore, flavonoids may play a role in the morphological appearance (short internodes and small, thick leaves) of light-demanding plants through their regulatory effect on auxin transport (Agati & Tattini, 2010).

Polysaccharides (8) are long chains of carbohydrates that play an important role in energy preservation, structural, and protective functions in plants (Bokov et al., 2020). As constituents of the cell wall, they are part of the plant’s first line of defense, acting as a physical barrier against pathogens (Vorwerk et al., 2004). This effect may contribute to the protection of medicinal plants on the kurgans from the increased risk of infection due to the proximity of arable fields, which are potential sources of plant diseases.

### 4.7 Soil potassium

We found that the potassium content of the soil positively influenced the presence of medicinal plants that produce different organic acids and their derivatives (such as aliphatic carboxylic acids: e.g., *Stenactis annua*, *Tragopogon dubius*; phenolic acids: e.g., *Cirsium arvense*, *Achillea collina*; and tannins: e.g., *Cirsium arvense*, *Rumex patientia*). This may be the result of a complex physicochemical and biological plant-soil feedback interactions that occur in the rhizosphere, and influence the potassium availability and the production of organic acids in plants (Sindhu et al., 2022).

Organic acids play an important role in plant-soil interactions by enhancing nutrient solubilization and attracting beneficial microorganisms (Sindhu et al., 2022). Organic acids enter the soil in the form of root exudates or through the decomposition of plant residues. Additionally, microorganism communities attracted by the released organic acids can also contribute to the production of further organic acids in the rhizosphere resulting in a positive feedback mechanism (Sindhu et al., 2022). Organic acids are crucial for the solubilization of otherwise unavailable potassium for uptake and utilization by plants. Adequate potassium sources maintain the transport of organic acids within plants and ensure their growth and development (Sindhu et al., 2022).

Due to the important role of potassium in water uptake and osmoregulation, its adequate supply improves the plant’s tolerance to drought stress (Mahiwal & Pandey, 2022). Greater drought tolerance is beneficial for medicinal plants on kurgans, as the elevated, hill-like shape of the mounds makes them the driest elements of the landscape, far from the water table and characterized by high levels of water runoff on their slopes.

Due to their ability to bind proteins and other macromolecules and to chelate metal ions, tannins enhance plant’s defense mechanisms against herbivores (as toxins and feeding deterrents) and pathogens (due to antimicrobial activity) and offer protection against heavy-metal toxins (Constabel et al., 2014). Phenolic acids are also involved in plant defense against ultraviolet radiation (due to antioxidant activity), pathogens (due to antimicrobial activity), and herbivores (as pest-feeding induced toxins) (Kessler & Kalske, 2018; Pant et al., 2021).

The higher phytochemical diversity observed on kurgans with higher potassium levels may be attributed to the multiple roles of potassium in plant physiological processes (Mahiwal & Pandey, 2022), such as cell metabolism, as well as the aforementioned plant-soil biological feedback interactions related to organic acid synthesis that enhance potassium availability to plants (Sindhu et al., 2022). Adequate potassium supply can contribute to increased production of secondary metabolites as part of the plant’s defense mechanisms (Mahiwal & Pandey, 2022).

## 5 CONCLUSION AND OUTLOOK

We revealed that ancient burial mounds (kurgans) can provide favorable microhabitats for a wide range of medicinal plants, including several typical grassland-related species. They can also act as repositories for a wide range of secondary metabolites produced by medicinal plants. Kurgans with different characteristics host different sets of medicinal plants, producing different secondary metabolite groups that assist species persistence by providing defense against herbivores, pathogens, UV radiation and drought. By producing metabolites, medicinal plants provide important ecosystem services through their healing and health-protective effects.

Although the collection of medicinal plants from the kurgans is not permitted due to their protected status, kurgans can serve as important hotspots and gene reserves for the conservation of medicinal plants, especially grassland-related and rare weed species, which are scarce in intensively used agricultural landscapes. Kurgans holding ancient or recovering grassland vegetation can serve as seed propagation sites, providing locally adapted ecotypes of medicinal plants for various restoration projects or for establishing sown parcels for medicinal plant production. They can also function as demonstration sites, introducing the typical local medicinal flora for local communities.

## ACKNOWLEDGEMENTS

The study was supported by the Hungarian Research, Development, and Innovation Office (grant numbers: NKFI FK 135329, KKP 144096); ÁB was supported by the NKFI FK 146506 project.

## AUTHOR CONTRIBUTIONS

Field data was collected by BD and OV, with contributions of ÁB. Field data preparation was performed by OV, BD and ÁB. Data on plant secondary metabolites and medicinal plants were collected by RE. Data analyses were performed by BD with contributions from AL. The first draft of the manuscript was written by RE and BD, and all authors commented on the manuscript. All authors contributed to the study conception and design, and read and approved the final manuscript.

## DATA AVIABILITY STATEMENT

Additional details and references on the classification of secondary metabolite, as well as on the secondary metabolite content and medicinal effects of the medicinal plants found on the studied kurgans, can be found in the supplementary material of this article.

