## Supporting information for "Small grassland patches are hotspots for medicinal plants and associated phytochemical diversity in European agricultural landscapes"

**Table S1.** Medicinal plants found on the studied kurgans, their main secondary metabolites and their medicinal properties and uses.

| | **Plant name** | **Main secondary metabolites** | **Therapeutic effects/potentials/uses** | **References** |
|---|---|---|---|---|
| 1 | *Abutilon theophrasti* Medik. | gallic acid, protocatechuic acid (trihydroxybenzoic acid), tannin, catechin (flavanol), vanillic acid, caffeic acid, ferulic acid, rutin, quercetin, syriacusin (naphthalenes and derivatives), mucilage, inositol and naphthoquinones, glucuronic acid, glucose, fructose, xylose, amino acid, fatty oil, sterols, free and glycosylated triterpenes, sugars and mixed polysaccharides (mucilages), oleanolic acid, triterpenoid sapogenin | expectorant, diuretic, analgesic, anti-inflammatory, anthelmintic, demulcent, aphrodisiac, emollient | Hassan et al., 2021 |
| 2 | *Achillea collina* J. Becker | essential oil (e.g. chamazulene), flavonoids | anti-inflammatory, antispasmodic, antiseptic, wound healing | Ph.Hg.VIII., 2004 |
| 3 | *Acinos arvensis* (Lam.) Dandy | Essential oil, tannin, flavonoids (lavones, flavonols and their glycosides), flavanone glycosides | antiseptics, stimulants, tonics, antispasmodics treat coughs, melancholy, toothache, sciatica, neuralgia, gastrointestinal disorders | Stojanovic et al., 2009 Jovanovic et al., 2005 Golubović et al., 2014 |
| 4 | *Agrimonia eupatoria* L. | tannins, flavonoids, phenolic acids | astringent, anti-inflammatory, digestive, diuretic, choleretic, antibiotic | EMA/HMPC/ 680597/2013, 2015 Paluch et al., 2020 |
| 5 | *Ailanthus altissima* (Mill.) Swingle | alkaloids, quassinoids, phenylpropanoids, triterpenoids, and essential oils | treat asthma, epilepsy, spermatorrhea, bleeding, ophthalmic disease | Li et al., 2021 |
| 6 | *Ajuga chamaepitys* (L.) Schreb. | iridoids, flavone glycosides, triterpenoids, neoclerodane diterpenoids, steroidal glucosides, phenylethanoids, flavonoids | wound healing, diuretic, tonic, emmenagogue, perspiration properties, menses remover | Venditti et al., 2016 Küpeli Akkol et al., 2020 |

| | | | | |
|---|---|---|---|---|
| 7 | *Allium scorodoprasum* L. | vanillic acid, protocatechuic acid, phydroxybenzoic acid, gallic acid, protocatechuic aldehyde, rutin, m-coumaric acid, p-coumaric acid, caffeic acid, quercetin, catechin, ferulic acid, naringenin, kaempferol, chlorogenic acid and rosmarinic acid, flavonoid, carotenoid, and chlorophyll | diabetes control and vision enhancement, antibacterial, antifungal, antioxidant, antiviral, diuretic, antibacterial, antifungal, antihypertensive, hepatoprotective, anti-obesity, antitumor | Ekici et al., 2022 |
| 8 | *Allium vineale* L. | flavonoids: hrysoeriol-7-O-[200-O-E-feruloyl]-b-D-glucoside, chrysoeriol , and isorhamnetin-3-<br>b-D-glucoside | antifungal | Demirtas et al., 2013 |
| 9 | *Alyssum alyssoides* L. | phenolic acids, flavonoids | treat male reproductive function disorders | Nejatbakhsh et al., 2016<br>Tsiftsoglou et al., 2019 |
| 10 | *Amaranthus retroflexus* L. | fatty acids, in particular linoleic and linolenic ones, essential amino acids, starch flavonoids (e.g. rutin and quercetin), alkaloids (e.g. amaranthine), sesquiterpenes, phenolic acids, and volatiles | treat cold, flu, hypercholesterolemia | Sargın et al., 2013<br>Fiorito et al., 2017 |
| 11 | *Ambrosia artemisiifolia* L. | sesquiterpene lactones | antiproliferative, cytotoxic, anti-inflammatory, antimicrobial | Chalchat et al., 2004<br>Solujic et al., 2008 , Kovács et al., 2022 |
| 12 | *Amorpha fruticosa* L. | isoflavonoids and their derivatives<br>called rotenoids, prenylated stilbenoids (phenolic compound), phenylpropanoids, volatile<br>terpenoids and fatty oils, geranyl-isoflavones | antidiabetic, anti-inflammatory, wound healing, antimicrobial, antibacterial, anti-tumor | Kozuharova et al., 2017 |
| 13 | *Anagallis arvensis* L. | saponins and flavonoids | wound healing | Lopez et al., 2011<br>Yasmeen et al., 2020 |
| 14 | *Anchusa officinalis* L. | phenolic compounds (caffeic acid, rosmarinic acid) | antioxidant, antibacterial, anti-inflammatory | Boskovic et al., 2018<br>Cartabia et al., 2021 |

| | | | | |
|---|---|---|---|---|
| 15 | *Anthemis arvensis* L. | sesquiterpene lactones | treat respiratory diseases, migraine, nose allergy, sneeze, hair loss, anxiety, insomnia, joints pains, pulmonary allergy, sore throat (pharyngitis) | Senouci et al., 2019 Vukovic et al., 2006 Vujisic et al., 2006 |
| 16 | *Anthriscus cerefolium* (L.) Hoffm. subsp. *trichosperma* (Spr.) Arc. | essential oil (methyl chavicol), flavone glycosides, fatty oil, bitter substances, minerals (Fe, Mg), vitamins (A, C) | circulatory improver, treat kidney-, bladder-, and digestive diseases | Bernáth, 2000 |
| 17 | *Arctium lappa* L. | inulin, mucilage, essential oil, polyacetylenes (arctinin, arctinol), polyphenols, lignans (arctiin), sterols | metabolism booster, treat liver-, gallbladder diseases, rheumatism, atopic dermatitis (eczema), anti-dandruff, hair loss | Bernáth, 2000<br><br>EMA/HMPC/ 246763/2009, 2010 |
| 18 | *Arenaria serpyllifolia* L. | Flavonoids and xanthones containing compounds such as epicatechin, japonicumone, quercetin etc. | treat bladder complaints such as acute and chronic inflammation of bladder, remove kidney lime stones, increase kidney function, fever, detoxify, improve eyesight, relief cough | Chandra & Rawat, 2015 |
| 19 | *Aristolochia clematitis* L. | aristolochic acids, astringent compounds, essential oil | treat atopic dermatitis (eczema), ulcerative wounds | Dános Béla, 2006 |
| 20 | *Artemisia absinthium* L. | absinthin, artabsin (sesquiterpene lactones type bitter compounds), essential oil | appetizer, digestive stimulant, choleretic, carminative | EMA/HMPC/ 751490/2016, 2020 |
| 21 | *Artemisia austriaca* Jacq. | Essential oil (camphor, 1,8-cineole, camphene, beta-fenchyl alcohol) | diuretic, deworming, fever anticandidal activity | Yigit et al., 2009 Razavi et al., 2014 |
| 22 | *Artemisia pontica* L. | sesquiterpene lactones | sedative, appetizer | Nigam et al., 2019 Todorova et al., 1996 |
| 23 | *Artemisia santonicum* L. | Essential oil | anthelmintic, treatment of diabetes | Kordali et al., 2005 |

| | | | | |
|---|---|---|---|---|
| 24 | *Artemisia vulgaris* L. | flavonoids, phenolic acid, organic acid, sesquiterpene, sesquiterpenic acid, sesquiterpene glucoside, sesquiterpene lactone, lignan glucoside, monoterpene, monoamine neurotransmitter, essential oil | stomach pain-relieving agent, treat gastric ulcers, hepatitis, convulsive crisis, neonatal jaundice, tonic, hypoglycemic, emmenagogue, anti-septic, expectorant, diuretic, analgesic, antihelminthic, treat rheumatism, asthma, cancer, epilepsy | Abiri et al., 2018 |
| 25 | *Asparagus officinalis* L. | steroid-saponins (asparagoside), amino acid (asparagine), flavonoids | diuretic | Dános Béla, 2006 |
| 26 | *Asperugo procumbens* L. | flavonoids, phenolics and tannins, fatty acids | strengthen the nervous system and the heart, counteract dementia, as an antispasmodic and tranquillizer, treat skin infections and herpes, liver diseases, asthma, improving digestion, stomach strengthening | Lazarski, 2021 |
| 27 | *Astragalus glycyphyllos* L. | saponins | antihypertensive, diuretic, anti-inflammatory, anti-tumor, | Georgieva et al., 2021 |
| 28 | *Atropa bella-donna* L. | amino acid (ornithine) tropane alkaloid (atropine, hyoscyamine) | pupil dilator | Ph.Hg.VIII., 2004 Kwakye et al., 2018 |
| 29 | *Avena sativa* L. | protein, minerals, lipids, β-glucan, a mixed-linkage polysaccharide, avenanthramides (hydroxycinnamic acid), an indole alkaloid-gramine, flavonoids, flavonolignans, triterpenoid saponins, sterols, and tocols | treat minor inflammations of the skin (such as sunburn), wound healing | Dános Béla, 2006 EMEA/HMPC /368600/2007, 2008 EMEA/HMPC /202966/2007, 2008 |
| 30 | *Ballota nigra* L. | phenylpropanoids, flavonoids, diterpene bitter compounds, iridoids, phenol carboxylic acids | treat flu, cough, induce neuro sedative activities, against anxiety | Ph.Hg.VIII., 2004 ESCOP & Roberta Hutchins, 2015 |

| | | | | |
|---|---|---|---|---|
| 31 | *Bryonia alba* L. | triterpenes (cucurbitacin glucosides), fatty acids (trihydroxyoctadecadienoic acids THODA), alkaloids (bryonicine), flavonoids (saponarin, vitexin, isovitexin 5, 7, 4'-trihydroxy flavone 8-C-glucopyranoside, lutonarin, isoorientin; glycosides, 22-deoxocucurbitosides A and B, 22-deoxocucurbitacin D), triterpenoids, (cucurbitacin L ), sterols (23, 24-dihydrocucurbitacin B, 23, 24,-dihydrocucurbitacin D), arvenin IV; lipids, proteins steroids, saponins, carbohydrates | treat oedema, anthelmintic, convulsions, headaches, bruises, pneumonia | Kujawska & Svanberg, 2019<br>Panossian et al., 1997 |
| 32 | *Calepina irregularis* (Asso) Thell. | glucosinolate (3-(methylsulfinyl) propyl isothiocyanate (degradation product of glucoiberin) and 3-(methylsulfonyl)propyl isothiocyanate (degradation product of glucocheirolin) | treat stomach ache | Rahman et al., 2018<br>Zekić et al., 2016 |
| 33 | *Camelina microcarpa* Andrz. | carbohydrates | hypoglycemic, hypolipidemic, antioxidant | Tsykalo & Trzhetsynskyi, 2021 |
| 34 | *Cannabis sativa* L. | cannabinoid Δ9-Tetrahydrocannabinol (THC),<br>Cannabidiol (CBD) | antibacterial, sedative, antispasmodic, analgesic, anti-inflammatory, narcotic | Bernáth, 2000<br>Ph. Eur. 11.5, 2024 |
| 35 | *Capsella bursa-pastoris* (L.) Medik. | flavonoids (diozmin), biogenic amines | acting on the uterus, reducing menstrual bleeding | EMA/HMPC/ 262766/2010, 2011 |

| | | | | |
|---|---|---|---|---|
| 36 | *Carex praecox* Schreb. | lignans, vanillic acid, flavonoids (tricin, quercetin), a chalcone (cilicinone B), stilbenoids (resveratrol, cis- and trans-ε-viniferin, cis-miyabenol C, kobophenol A, carexinol A),lignan derivative, phenolic compounds (phenolic acids, phenylpropanoids) | treat stomach and bowel disorders, amenorrhea, bronchitis, hematopoietic disorders, tumors, infectious diseases, pain and fever, diabetes, skin diseases, problems concerning the circulation, digestive, respiratory and reproductive organs | Dávid et al., 2022<br>David et al., 2021 |
| 37 | *Carlina vulgaris* L. | polyphenols (chlorogenic acid), minerals | antioxidant properties | Strzemski et al., 2017 |
| 38 | *Caucalis platycarpos* L. | flavonoids and phenolic acids | antitumor activity | Plazonic et al., 2011 |
| 39 | *Celtis occidentalis* L. | tyramine and octopamine derivatives, flavonoids, phenolic acids, anthocyanins, triterpenes, lignans and amide derivatives | treat sore throat and aid during menstruation, treating jaundice | Ayanlowo et al., 2020 |
| 40 | *Centaurea cyanus* L. | polysaccharide | treat minor ocular inflammation, anti-inflammatory effects | Garbacki et al., 1999 |
| 41 | *Centaurea pannonica* (Heuff.) Simk. | essential oil | antibacterial effect | Csupor et al., 2011<br>Milošević Ifantis et al., 2013<br>Formisano et al., 2010 |
| 42 | *Centaurea solstitialis* L. | sesquiterpene lactones (solstitialin)<br>neurotoxic amino acids | antipyretic, analgesic, antitumor activity | Csupor et al., 2011 |
| 43 | *Chenopodium album* L. | flavonoid as phenolic, amide, saponin, cinnamic acid amide, alkaloid, phenols and lignans, proteins, calcium and vitamins A, carotenoids, vitamin C, iron content | anthelmintic, cardiotonic, carminative, digestive, diuretic and laxative | Poonia & Upadhayay, 2015 |
| 44 | *Chenopodium hybridum* L. | flavonoids and phenolic acids | analgesic agent | Podolak et al., 2016 |
| 45 | *Cichorium intybus* L. | inulin, cichoric acid, cichoriin, sesquiterpene lactones, diterpenes | choleretic, digestive disorders, | EMA/HMPC/121816/2010, 2013 |

| | | | | |
|---|---|---|---|---|
| 46 | *Cirsium arvense* (L.) Scop. | flavonoid compounds, phenolic acids, tannins, sterols and triterpenes | diuretic, astringent, anti-inflammatory | Nazaruk, 2008 |
| 47 | *Cirsium canum* (L.) All. | chlorogenic acid, caffeic acid, p-coumaric acid, protocatechuic acid, p-hydroxybenzoic acid, vanillic acid, syringic acid, trans-cinnamic acid, luteolin-7-glucoside, apigenin-7-glucoside, kaempferol-3-glucoside, linarin, apigenin, rutoside, luteolin and kaempferol | antimicrobial | Kozyra et al., 2015 |
| 48 | *Cirsium vulgare* (Savi) Ten. | flavonoid compounds, phenolic acids, tannins, sterols and triterpenes | anxiolytic activity | Nazaruk, 2008 |
| 49 | *Consolida orientalis* (J. Gay) Schrödinger | diterpene alkaloids, anthocyanins, flavonoids | antihypertensive, laxative, vasodilator | Bernáth, 2000 |
| 50 | *Consolida regalis* S. F. Gray | diterpene alkaloids, anthocyanins, flavonoids | antihypertensive, laxative, vasodilator | Bernáth, 2000 Dános Béla, 2006 |
| 51 | *Convolvulus arvensis* L. | carbohydrates, coumarins, saponins, flavonoids, lipids, steroids or terpenoids, sugar derivatives of kaempferol and quercetin, tannins, alkaloids, lactones, proteins or amino acids, arvensic acids A-D (resin glycoside), and vitamin E | antispasmodic, anti-inflammatory, anti-swelling, treat painful joints, treat flu, haemostatic, antiangiogenic, laxative | Salehi et al., 2020 |
| 52 | *Conyza canadensis* (L.) Cronquist | polysaccharides and their conjugates with proteins | treat cough, cold and asthma diuretic, haemostatic, tonic, antidiarrheal, antifungal, antimicrobial, anthelmintic, astringent, antioxidant, anti-inflammatory, anti-coagulant, anticancer, mutagenic, gastric protective, and skin de-pigmentation effect | Šutovská et al., 2022 |

| | | | | |
|---|---|---|---|---|
| 53 | *Coronilla varia* L. | cardenolide (hyrcanoside) coumarins (daphnoretin, scopoletin, umbelliferone) | cytotoxic and antileukemic activity | Williams & Cassady, 1976 TOPPEL et al., 1987 |
| 54 | *Crataegus monogyna* Jacq. | procyanidins, flavonoids, caffeic acid, chlorogenic acid | antihypertensive, cardiac failure | Ph.Hg.VIII., 2004 |
| 55 | *Crepis biennis* L. | Sesquiterpens (ixerin F in roots) flavonoids (luteolin , luteolin 7-O-glucoside , luteolin 7-O-glucuronide (81), luteolin 7-O-gentiobioside) chlorogenic acid, caffeoyltartaric acid, 3,5-di-O-dicaffeoylquinic acid , cichoric acid fatty acid (in seed, vernolic acid, crepenynic acid, palmitic acid, stearic acid, oleic acid, linoleic acid) | anti-inflammatory, antioxidant, digestive | Badalamenti et al., 2022 |
| 56 | *Cuscuta campestris* Yuncker | flavonoids (hyperoside and quercetin), polysaccharides, alkaloids | hepatoprotective, stimulating effect on lymphocyte proliferation and antibody production, anti-inflammatory and anti-proliferative activities | Lee et al., 2011 |
| 57 | *Cynodon dactylon* (L.) Pers. | saponins | diuretic | Bernáth, 2000 Shendye & Gurav, 2014 |
| 58 | *Cynoglossum officinale* L. | pyrrolizidine alkaloids (heliosupine, rinderine, echinatine, 7-angelolyl heliotridine | antihemorrhagic, antiseptic, diuretic, treat venereal diseases | Joshi, 2016 |
| 59 | *Datura stramonium* L. | tropane alkaloids (hyoscyamine, scopolamine, atropine) | asthma, antispasmodic | Ph.Hg.VIII., 2004 |

| | | | | |
|---|---|---|---|---|
| 60 | *Descurainia sophia* (L.) Webb | fatty acid, monoterpenes, sesquiterpenes, essential oil, cardiac glycosides, flavonoids, phenols, nor-lignans, descurainoside (sulfur glycoside), flavonoids, coumarin | anthelmintic, antioxidant, radical scavenging, analgesic, anti-inflammatory, antipyretic effects | Nimrouzi & Zarshenas, 2016<br>Hsieh et al., 2020 |
| 61 | *Dipsacus laciniatus* L. | iridoid glycosides, phenolic glucoside (dipsaicin), ursolic acid | treat respiratory diseases, | Papp et al., 2011 |
| 62 | *Echium vulgare* L. | phenolic acids, flavonoids, pyrrolizidine alkaloid, naphthoquinones sterone, polysaccharides, unsaturated fatty acids, allantoin; tetracarboxylic acids (citric acid, fumaric acid, malic acid and succinic acid); lignans; tannins, uridine; saponin | treat respiratory diseases, cough | Wang et al., 2022 |
| 63 | *Elaeagnus angustifolia* L. | alkaloids, flavonoids, saponins, tannins, flavonoids | diuretic, wound healing, anti-inflammation, antipyretic, treat cough, urinary tract infection, muscle strain | Hosseinzadeh et al., 2003 |
| 64 | *Elymus repens* (L.) Gould | tricin (flavone, poly-fructosan), mucilage (polysaccharide), essential oil, silicon dioxide | diuretic, treat tracheitis | Ph.Hg.VIII., 2004 |
| 65 | *Epilobium tetragonum* L. | sterols, triterpenes, fatty acids, macrocyclic tannins, flavonoid glycosides | inhibit proliferation of human prostate cells | Vitalone et al., 2003 |
| 66 | *Equisetum arvense* L. | minerals (silicon dioxide), flavonoids | diuretic, topical treatment for joint disorders | Ph.Hg.VIII., 2004 |
| 67 | *Erigeron acer* | triterpene (alpha-amyrin, beta-amyrin), caffeic acid, quercetin, 4'-hydroxywogonin-7-O-beta-D-glucuronic acid glycoside | treat tooth-ache, bruises, arthritis | Pieroni et al., 2004<br>Yan et al., 2008 |
| 68 | *Erodium ciconium* (L.) L'Hérit. | alkaloids, tannins and flavonoids, essential oil, phenolic compounds | astringent, antiseptic | Fecka & Cisowski, 2002 |

| | | | | |
|---|---|---|---|---|
| 69 | *Erodium cicutarium* (L.) L'Hérit. | gallic acid, protocatechuic acid, 3-O-galloylshikimic acid, 3-O-(6”-O-galloyl)-β-D-galactopyranoside, corilagin, didehydrogeraniin (dehydrogeraniin), geraniin, hyperin, isoquercitrin, methyl gallate 3-O-β-D-glucopyranoside, and rutin phenolic compounds, essential oil, tannin, alkaloid | treat dermatological diseases, respiratory diseases, influenza, heart problems, stomach ache, low and high blood pressure, antihemorrhagic, wound-healing, anti-inflammatory, antimicrobial and antiviral activities | Munekata et al., 2019 |
| 70 | *Eryngium campestre* L. | triterpene saponins, essential oil | treat tracheitis, bronchitis | Dános Béla, 2006 |
| 71 | *Erysimum diffusum* Ehrh. | cardiac glycosides, astringent compounds, vitamin C | treat heart failure, increase myocardial contraction | Dános Béla, 2006 |
| 72 | *Euphorbia cyparissias* L. | diterpenes (cyparissins A, B) | cytotoxic activity against A2780 human ovarian cancer cells | Lanzotti et al., 2015 |
| 73 | *Euphorbia esula* L. | diterpenes (presegetane, jatrophane, paraliane, isopimarane, euphorbesulins) | cytotoxic, antibacterial, anti-inflammation, treat warts, cancer, swelling | Amtaghri et al., 2022 |
| 74 | *Euphorbia helioscopia* L. | jatrophane-type diterpenoids | antifungal, antibacterial, antidiarrheal, diuretic, treat tuberculosis, scabies, cancer | Amtaghri et al., 2022 |
| 75 | *Euphorbia virgata* W. et K. | diterpenoids | antimicrobial, antifungal, wound-healing, treat eczema | Amtaghri et al., 2022 |
| 76 | *Fallopia convolvulus* (L.) A. Löve | flavonoids | anticancer properties | Kim et al., 2000<br>Olaru et al., 2015 |
| 77 | *Fraxinus angustifolia* Vahl subsp. *pannonica* Soó et Simon | coumarins (fraxoside, hydrolyzable fraxetol, monosaccharide (mannitol), organic acids, tannins, sugars. In bark infusions, glucose, fraxin, tannins, bitter principles<br>coumarins, secoiridoids, phenylethanoids, lignans, flavonoids, phenolic compounds | treat inflammatory diseases: rheumatism, arthritis, gout leaves: antidiarrheal, anthelmintic, bark: against gallstones, passive hemorrhages, gout, cholelithiasis, antipyretic | Ayouni et al., 2016 |

| | | | | |
|---|---|---|---|---|
| 78 | *Fumaria officinalis* L. | benzylisoquinoline alkaloids, fumarin, resin, amino acid, bitter compounds, mucilage, caffeic acid, flavonoids | choleretic, bile-regulating, smooth muscle relaxant | Ph.Hg.VIII., 2004<br><br>EMA/HMPC/ 574766/2010, 2011 |
| 79 | *Galium aparine* L. | phenols, tannins, alkaloids, anthraquinones, coumarins, iridoids asperuloside, alkanes, flavonoids, saponins | antimicrobial, anticancer, hepatoprotective, sedative | Al-Snafi, 2018 Dános Béla, 2006 |
| 80 | *Galium spurium* L. | phenolic acids, flavonoids, and iridoids | anticonvulsant: treat epilepsy, treat bones, sinews pain, anticancer | Orhan et al., 2012 |
| 81 | *Galium verum* L. | tannins, triterpenes, essential oil,<br>organic acids, carotenoids, flavonoids, iridoid glycosides, anthracene derivatives | anti-inflammatory, antioxidant, antiseptic, restorative, choleretic, analgesic, diuretic, antispasmodic, estrogenic, anticancer, treat kidney, liver and respiratory diseases | Dános Béla, 2006 Zaichikova et al., 2020 |
| 82 | *Geum urbanum* L. | gein (eugenol glycoside), essential oil, bitter compounds, astringent compounds | anti-inflammatory: inflammation of the oral mucosa, gingivitis, gastrointestinal and biliary disorders | Bernáth, 2000 Dános Béla, 2006 |
| 83 | *Glechoma hederacea* L. s.str. | diterpenes, iridoids, essential oil | expectorant, mild sedative | Bernáth, 2000 Dános Béla, 2006 |
| 84 | *Gleditsia triacanthos* L. | triterpenes, sterols, flavonoids, alkaloid, phenolics and their derivatives | pupil dilator, analgesic, anti-HIV activity | J. Zhang et al., 2016 |
| 85 | *Helianthus annuus* L. | fatty acid (linoleic acid, oleic acid) proteins, sterol | ointment base | Ph.Hg.VIII., 2004 |
| 86 | *Hieracium pilosella* L. | phenolic compounds, flavonoids, coumarins, sesquiterpene lactones, terpenoids, phytosterols | diuretic, urinary tract disorders | EMA/HMPC/ 680374/2013, 2015<br>Willer et al., 2021 |
| 87 | *Hyoscyamus niger* L. | tropane alkaloid (atropine, hyoscyamine, scopolamine) | treat asthma, rheumatic pain, neuralgia | Ph.Hg.VIII., 2004 |

| | | | | |
|---|---|---|---|---|
| 88 | *Hypericum perforatum* L. | naphtodianthrone hypericin (hypericin, pseudo-hypericin), phloroglucinol derivative (hyperforin, adhyperforin) flavonoids (hyperoside), essential oil, astringent compounds | sedative, antidepressant, antianxiety, anti-inflammatory, stomach ulcer prevention, liver-protective, treat slow-healing burns | Ph.Hg.VIII., 2004 |
| 89 | *Inula britannica* L. | sesquiterpene-lactones (guaianolids, eudesmanolids, germacranolids, xanthanolides), flavonoids, essential oils, triterpenoids, diterpenes | treat edemas, reduction in nausea, prevention of vomiting, treat cold-related coughs | Seca et al., 2014<br>Yang et al., 2021 |
| 90 | *Juglans regia* L. | naphthoquinone derivative (juglone), flavonoids, astringent compounds, essential oil, vitamin C | treat skin disorders and minor wounds, treat enteritis | EMA/HMPC/ 346737/2011, 2013<br>Bernáth, 2000 |
| 91 | *Lamium purpureum* L. | iridoids, essential oil | tonics, treat constipation, anti-inflammatory, analgesic | Akkol et al., 2008 |
| 92 | *Leonurus cardiaca* L. | iridoids (leonuride ajugoside, galiridoside), bitter diterpenoids (leocardin , leosibiricin), triterpenes (ursolic acid , oleanolic acid, corosolic acid, euscaphic acid, ilelatifol D) pyrrolidine alkaloids (stachydrine, betonicine, turicin, imines guanidine derivative (leonurine = syringic acid ester and 4-uanidino-1-butanol amine) choline flavonoids phenolic compounds phenylpropanoid glycosides (lavandulifolioside)<br>essential oils, sterols (b-sitosterol and stigmasterol) tannins | treat cardiac anxiety, circulatory disorder, antihypertensive | Wojtyniak et al., 2013<br>Dános Béla, 2006 |
| 93 | *Linaria vulgaris* Mill. | flavonoids (linarin), aurone glycoside (aureusin), alkaloid | liver and biliary disorders, laxative, diuretic | Dános Béla, 2006 |

| | | | | |
|---|---|---|---|---|
| 94 | *Lotus corniculatus* L. | Phenolics, flavonoid, kaempferitrin, saponin derivatives, oleanolic acid, b-sitosterol, | carminative, antipyretic, antispasmodic, sedative, anti-cancer, anti-inflammatory | Yerlikaya et al., 2019<br>Zhao et al., 2019<br>Koelzer et al., 2009<br>Rafiq et al., 2013<br>Foo et al., 1996 |
| 95 | *Lycium barbarum* L. | polysaccharides, carotenoids and related compounds, flavonoids (rutin, gentisic acid, and quercetin), betaine (amino acid derivatives), cerebroside, beta-sitosterol, p-coumaric acid, vitamins, minerals | metabolism booster, cardiovascular benefits, diabetes, immune modulation, Interleukin 12 (IL-12), phagocytic action, anti-inflammation: skin protection from UV radiation, anti-cancer, cytoprotectant | Amagase & Farnsworth, 2011<br>Ph.Hg.VIII., 2004 |
| 96 | *Lycopus europaeus* L. | caffeic acid derivatives (lithospermic acid), astringent compounds, flavonoids, essential oil, diterpenes | treat hyperthyroidism, premenstrual syndrome | Dános Béla, 2006 |
| 97 | *Lysimachia nummularia* L. | flavonoids (kaempferol, quercetin myricetin aglycones), triterpene saponins | anti-inflammatory | Toth et al., 2016 |
| 98 | *Malva neglecta* Wallr. | flavonoids, phenolic acids, dimercaptan, phenylpropanoids (isoeugenol), amino sulfonic acid<br>(taurine), sesquiterpenoid (patchoulane), fatty acid (isopropyl myristate, methyl 12-methyltetradecanoate, oleic acid), colchicine | treat cough and cold gastrointestinal disorders, wound-healing | Saleem et al., 2020<br>Ph.Hg.VIII., 2004<br>EMA/HMPC/ 749510/2016, 2018 |
| 99 | Malva sylvestris L. | mucilage compounds, astringent compounds, anthocyanidin (malvidin-glucoside), flavonoids | expectorant, cough suppressant, treat eczema | Ph.Hg.VIII., 2004<br>Bernáth, 2000<br>Csupor & Szendrei, 2012 |
| 100 | *Marrubium peregrinum* L. | diterpene bitter compound (marrubiin), astringent compounds, essential oil | choleretic, mucolytic | Dános Béla, 2006 |
| 101 | *Matricaria chamomilla* L. | essential oil (matricin) flavonoids, coumarins, mucilage compounds | antispasmodic, anti-inflammatory, antiseptic | Ph.Hg.VIII., 2004 |

| | | | | |
|---|---|---|---|---|
| 102 | *Medicago sativa* L. | flavonoids, proteins, vitamins | diuretic, anti-asthmatic, anti-arthritic, anti-diabetic | Mansourzadeh et al., 2022 |
| 103 | *Melilotus officinalis* (L.) Pall. | Coumarin derivatives (melilotin), flavonoids | treat skin disorders and minor wounds, circulatory disorders, gastrointestinal and respiratory diseases, inflammation of varicose veins | Ph.Hg.VIII., 2004<br><br>EMA/HMPC/ 44166/2016, 2017<br>Dános Béla, 2006 |
| 104 | *Muscari comosum* (L.) Mill. | phenolic acids, flavonoids | antioxidant, anti-inflammatory, reduce post-prandial hyperglycemia | Larocca et al., 2018 |
| 105 | *Nigella arvensis* L. | flavonoids, phenolic acids, phytosterols, triterpenes, alfa-tocopherol, beta-carotene, essential oils, fatty acids (linoleic<br>acid, oleic acid) | anti-inflammatory, enhance male potency, improve memory, antioxidant, anti-cancer, treat psoriasis | Salehi et al., 2021 |
| 106 | *Oenothera biennis* L. | fatty acid (gamma linolenic acid) | treat cardiovascular diseases, cosmetology | Ph.Hg.VIII., 2004<br>Csupor & Szendrei, 2012 |
| 107 | *Ononis arvensis* L. | phytosterols, triterpenes, lectins, flavonoids, (isoflavonoid glucoside, dihydroisoflavonoid), hydroxycinnamic<br>acids, coumarin (scopoletin, scopolin) | treat infections of<br>urinary tract, skin diseases | Gampe et al., 2019 |
| 108 | *Ononis spinosa* L. | triterpene saponins (onocerin), isoflavone (ononin, onospin, trifolirizin), flavonol (rutin, kaemferol), essential oil, astringent compounds, fatty oil, sterol | diuretic | Bernáth, 2000<br>Ph.Hg.VIII., 2004 |
| 109 | *Onopordum acanthium* L. | flavonoids, phenylpropanoids, lignans, triterpenoids, sesquiterpene lactones, sterols | anti-inflammatory, anticancer, cardiotonic | Csupor-Loffler et al., 2014<br>Garsiya et al., 2019 |
| 110 | *Oxalis europaea* Jord. | oxalic acid (organic acid), glycosides, phenols, flavonoids, tannins, | antipyretic, treat indigestion, anti-inflammatory, appetizer, treat mouth irritations, nausea, stomach cramps, antihemorrhagic, treat blood toxicity | Dzinyela et al., 2021 |

| | | | | |
|---|---|---|---|---|
| 111 | *Papaver rhoeas* L. | Alkaloids (rhoeadine, rhoeadic acid), mucilage compounds, anthocyanins | anti-inflammatory, mild cough suppressant, eye wash, throat gargle | Ph.Hg.VIII., 2004<br>Dános Béla, 2006<br>Bernáth, 2000 |
| 112 | *Phalaris canariensis* L. | polyunsaturated fatty acids (linoleic acid), sterols contents | treat: diabetes mellitus, hypertension, hypercholesterolernia | Ben Salah et al., 2018 |
| 113 | *Phlomis tuberosa* L. | iridoid glycosides | wound healing,<br>treat lung and throat diseases | Mukemre et al., 2015<br>Olennikov & Chirikova, 2017 |
| 114 | *Pimpinella saxifraga* L. | essential oil, coumarin derivatives | treat respiratory tract inflammation | Morlock & Lapin, 2018<br>Dános Béla, 2006 |
| 115 | *Plantago lanceolata* L. | iridoid glycoside (aucubin, catalpol), polyphenols, mucilage compounds, vitamin C, astringent compounds, caffeic acid derivatives, silicium dioxide | anti-inflammatory, treat pharyngitis and bronchitis, wound healing | Ph.Hg.VIII., 2004<br>Dános Béla, 2006<br>Bernáth, 2000<br>Csupor & Szendrei, 2012 |
| 116 | *Plantago major* L. | iridoid glycoside (aucubin, catalpol), polyphenols, mucilage compounds, vitamin C, phenol derivatives, bioflavonoids, tannic acid derivatives | anti-inflammatory, treat pharyngitis and bronchitis, wound healing | Dános Béla, 2006<br>Bernáth, 2000<br>Turgumbayeva et al., 2022 |
| 117 | *Podospermum canum* C. A. Mey. | flavonoids | anti-inflammatory, analgesic | Akkol et al., 2012 |
| 118 | *Polygonum aviculare* L. s.str. | flavonoids, astringent compounds, silicium dioxide | astringent, antibiotic, antihemorrhagic, diuretic, treat gastric ulcers, respiratory inflammation | Ph.Hg.VIII., 2004<br>Bernáth, 2000 |
| 119 | *Polygonum lapathifolium* L. | phenylpropanoids and their | anticancer, anti-tumor, antioxidative, anti-inflammatory, analgesic, antibacterial | Hailemariam et al., 2018 |

| | | | | |
|---|---|---|---|---|
| 120 | *Populus alba* L. | salicylic alcohol (saligenin), salicinoids (salicin, salicortin), flavonoids | expectorant, diuretic, anti-rheumatic, depurative, antiseptic, antipyretic, treat tooth decay, dysuria, eczema, diabetes, high blood pressure, asthma | Pobłocka-Olech et al., 2021 |
| 121 | *Populus* x *canadensis* Mönch | essential oil (humulene, caryophyllene) flavone glycosides (salicin, populin) astringent compounds, cinnamic acid derivatives, flavonoids | antibiotic, treat gout, hemorrhoid | Csupor & Szendrei, 2012 |
| 122 | *Portulaca oleracea* L. | alkaloids, terpenoid, organic acids, fatty acids, flavonoids, minerals, vitamins | treat gastrointestinal diseases, respiratory problems, liver inflammation, kidneys and bladder ulcers, insomnia, headaches, antipyretic, anti-inflammatory | Iranshahy et al., 2017 |
| 123 | *Potentilla reptans* L. | flavonoids | antidiarrheal, anti-inflammatory, treat tooth ache, dental problems, ulcers, sore throat, mastitis, hemorrhoids | Dános Béla, 2006<br>Augustynowicz et al., 2021<br>Tomczyk & Latté, 2009 |
| 124 | *Prunella vulgaris* L. | triterpenoids, phenolic acids, flavonoids, polysaccharides, sterols, phenylpropanoids, fatty acids, essential oils | antiviral, antibacterial, anti-inflammatory, immunomodulatory, anti-oxidative, anti-tumor, antihypertensive and hypoglycemic function | Ph.Hg.VIII., 2004<br>Pan et al., 2022<br>Bai et al., 2016 |
| 125 | *Prunus cerasifera* Ehrh. | polyphenolics, anthocyanins, carotenoids, flavonoids, organic acids, aromatic compounds, tannins, minerals, vitamins, antioxidant compounds | antibacterial, antifungal | Liu et al., 2020 |
| 126 | *Prunus spinosa* L. | flavonoids, procyanidins | gentle laxative, diuretic | Dános Béla, 2006<br>Bernáth, 2000 |
| 127 | *Ranunculus arvensis* L. | flavonoids, phenolics | anti-rheumatic, treat dermatological disorders: wounds, burns, psoriasis | Polat, 2016 |

| | | | | |
|---|---|---|---|---|
| 128 | *Reseda lutea* L. | benzyl isothiocyanate | antibacterial, anti-inflammatory, anti-HIV, anti-tumor | Radulovic et al., 2014 |
| 129 | *Robinia pseudoacacia* L. | Flavonoids (in flower) toxic proteins (in bark, robin, phasin) | gentle laxative, treat hyperacidity | Dános Béla, 2006 |
| 130 | *Rosa canina* L. s.str. | Vitamin C, B1, B2, P, carotenoids, carbohydrates, pectin, organic acids | vitamin supplement | Ph.Hg.VIII., 2004<br>Dános Béla, 2006<br>Bernáth, 2000 |
| 131 | *Rosa rubiginosa* L. s.str. | Vitamin C, B1, B2, P, carotenoids, carbohydrates, pectin, organic acids | vitamin supplement | Dános Béla, 2006<br>Bernáth, 2000 |
| 132 | *Rubus caesius* L. | astringent compounds (gallotannins, organic acids, flavonoids, | astringent, mild antispasmodic | Bernáth, 2000 |
| 133 | *Rumex acetosa* L. | flavonoids, astringent compounds, anthraquinones | root: enteritis, gastritis, gentle laxative; fruit: antidiarrheal, antibacterial, antifungal, treat psoriasis | Bernáth, 2000<br>Dános Béla, 2006<br>Vasas et al., 2015 |
| 134 | *Rumex acetosella* L. | anthraquinones (aloe-emodin, chrysophanol, physcion, emodin, sennoside A, B), naphthalene, stilbenoids (polyphenol), steroids, flavonoids, leucoanthocyanidins and phenolic acids | treat kidney disorders, swellings, gentle laxative (root), antidiarrheal (fruit), analgesic, diuretic, treat skin disorders (sores, rashes, wounds, ringworm), astringent | Vasas et al., 2015 |
| 135 | *Rumex crispus* L. | flavonoids, astringent compounds, anthraquinones (in root) | treat skin disorders (sores, rashes, wounds, ringworm), astringent | Dános Béla, 2006<br>Bernáth, 2000<br>Vasas et al., 2015 |
| 136 | *Rumex obtusifolius* L. | flavonoids, polyphenols, astringent compounds, anthraquinones (in root) naphthalene-diolester, tannin | root: enteritis, gastritis, gentle laxative; fruit: antidiarrheal, antibacterial, antifungal, treat psoriasis | Bernáth, 2000 |

| | | | | |
|---|---|---|---|---|
| 137 | *Rumex patientia* L. | flavonoids, polyphenols, astringent compounds, anthraquinones naphthalene-diolester, tannin (in root) | root: enteritis, gastritis, gentle laxative; fruit: antidiarrheal, antibacterial, antifungal, treat psoriasis | Bernáth, 2000<br>Dános Béla, 2006<br>Vasas et al., 2015 |
| 138 | *Rumex stenophyllus* Ledeb. | Anthraquinones, naphthalene-1,8-diols, flavonoids and stilbenoids | treat cough | Vasas et al., 2015 |
| 139 | *Salvia austriaca* Jacq. | diterpenoids, 7a-acetox-yroyleanone, 7a-hydroxyroyleanone flavonoids, tannins, triterpenes, essential oils, phenolic acids | antibacterial, antiviral, anti-inflammatory, antioxidant, treat cardiovascular disorders | Nagy et al., 1999<br>Janicsak et al., 2011 |
| 140 | *Salvia nemorosa* L. | flavonoids, tannins, triterpenes, essential oils, diterpenoids, phenolic acids | antibacterial, antiviral, anti-inflammatory, antioxidant, treat cardiovascular disorders | Chizzola, 2012 |
| 141 | *Salvia pratensis* L. | flavonoids, tannins and triterpenes, essential oils, diterpenoids, phenolic acids | antibacterial, antiviral, anti-inflammatory, antioxidant, treat cardiovascular disorders abdominal pains, skin diseases, ulcers | Srećković et al., 2022<br>Janicsak et al., 2011 |
| 142 | *Sambucus nigra* L. | flavonoids, saponins, chlorogenic acid, cyanogenic glycosides, essential oil, mucilage compounds | sudorific, diuretic | Ph.Hg.VIII., 2004<br>Bernáth, 2000 |
| 143 | *Scrophularia nodosa* L. | Iridoids, iridoid glycosides | wound healing | Stevenson et al., 2002 |
| 144 | *Secale cereale* L. | seed: starch, proteins, vitamin B<br>pollen: sterols, amino acids, fatty acids, minerals, vitamins | restorative, immunomodulator, adjuvant in prostate diseases. | Dános Béla, 2006<br>Kulichová et al., 2019 |
| 145 | *Senecio vernalis* W. et K. | pyrrolizidine alkaloids (senecivernine, senecionine, seneciphylline, integerrimine) | antidiarrheal, diuretic, emmenagogue, galactagogue | Seremet et al., 2018<br>Ahmadi-Noorbakhsh et al., 2011 |

| | | | | |
|---|---|---|---|---|
| 146 | *Silene vulgaris* (Mönch) Garcke | essential oils, phytoecdysteroids | immunomodulatory, treat liver diseases | Tuttolomondo et al., 2014 Rezaeieh et al., 2016 Sidana et al., 2017 |
| 147 | *Solidago gigantea* Ait. subsp. *serotina* (Ait.) McNeill | flavonoids, triterpene saponins, essential oil, astringent compounds, bitter compounds | diuretic, anti-inflammatory, immunomodulatory, antifungal, astringent, treat gout, effect on urinary tract, anti-rheumatism, blood purifier | Ph.Hg.VIII., 2004 Dános Béla, 2006 Bernáth, 2000 |
| 148 | *Sonchus oleraceus* L. | taraxa sterol (pentacyclic triterpenoid), flavonoids (apigenin 7-glucuronide, luteolin 7-glucoside), alkaloids, coumarins, saponins | analgesic, treat stomachic pain, headache, toothaches, hepatitis, anti-inflammatory, anti-rheumatism, depurative, diuretic, laxative, facilitate hepatic and intestinal function, general tonic | Sansanelli et al., 2017 Vilela et al., 2009 |
| 149 | *Stachys annua* (L.) L. | bitter compounds, astringent compounds | treat kidney and bladder disorders, respiratory diseases | Dános Béla, 2006 Kanjevac et al., 2023 |
| 150 | *Stachys germanica* L. | flavonoids, irridoids | treat gastric pain, painful menstruation | Kanjevac et al., 2023 Haznagy Radnai et al., 2006 |
| 151 | *Stellaria media* (L.) Vill. | saponins, minerals, essential oil | diuretic | Bernáth, 2000 Dános Béla, 2006 |
| 152 | *Stenactis annua* (L.) Nees | Terpenoids, sterols, flavonoids, organic acid | treat indigestion, enteritis, epidemic hepatitis, lymphadenitis, hematuria, acute inflammation, malaria obesity | L. Zhang et al., 2020 |
| 153 | *Symphytum officinale* L. | allantoin, astringent compounds, mucilage compounds, pyrrolizidine alkaloids | tissue regenerative, epithelializing, anti-inflammatory, treat muscle strain, joint disorders | EMA/HMPC/572846/2009, 2015 Bernáth, 2000 Dános Béla, 2006 |
| 154 | *Syringa vulgaris* L. | iridoids, seco-iridoids, lignans, phenylethanoid glycosides, phenolic compounds, essential oils | antipyretic, treat cold, cough, gastrointestinal disorders, wound-healing, anti-inflammatory | Filipek et al., 2019 |

| | | | | |
|---|---|---|---|---|
| 155 | *Tanacetum vulgare* L. | essential oil (thujone, camphor, borneol, umbellulone, artemisia ketone), parthenolide sesquiterpene lactones, polyacetylene, organic acid | anthelmintic, anti-rheumatic, treat inflammation of varicose veins | Bernáth, 2000 |
| 156 | *Taraxacum officinale* Weber | sesquiterpene lactone bitter compounds, sterols (taraxasterol), inulin, triterpenes, vitamin C, A, B | diuretic, choleretic | Ph.Hg.VIII., 2004<br>Bernáth, 2000<br>Dános Béla, 2006 |
| 157 | *Thlaspi arvense* L. | unsaturated fatty acids (erucic acid, linoleic acid), triacylglycerols | treat painful urination | Ballabh et al., 2008 |
| 158 | *Tragopogon dubius* Scop. | Flavonoids, phenylmethane derivatives (benzoic acid), phenylpropane derivatives (p-coumaric acid, caffeic acid) | treat stomach ache, diuretic, hypotensive, antirheumatic, antidiabetic, wound-healing | Abdalla & Zidorn, 2020 |
| 159 | *Trifolium pratense* L. | isoflavone (daidzein, genistein, formononetin, biochanin A) | sedative, analgesic (cancer, rheumatism, gout), treat whooping cough, gynecological problems | Nissan et al., 2007 |
| 160 | *Urtica dioica* L. | vitamins, triterpenes, sterols, flavonoids, amines, minerals, root: sterols, coumarins, astringent compounds, phenylpropane derivatives, lignan | diuretic, blood purifier, anti-inflammatory | Ph.Hg.VIII., 2004<br>Bernáth, 2000<br>Dános Béla, 2006 |
| 161 | *Verbascum phlomoides* L. | mucilage compounds, triterpene saponins, terpenoids, sugars, flavonoids, carotenoids, iridoids, caffeic acid derivatives, essential oil | treat upper respiratory diseases, gastrointestinal complaints, expectorant, cough suppressant, sudorific, anti-rheumatism, shampoo | Bernáth, 2000<br>Csupor & Szendrei, 2012 |
| 162 | *Verbena officinalis* L. | iridoid-glycoside (verbenin, verbenalin), caffeic acid derivatives, astringent compounds, saponins, flavonoids | anti-inflammatory, gentle analgesic, inhibits increased thyroid activity, treat menstrual disorders, galactagogue, metabolism booster, treat bruise, frostbite, diaper rash, burn | Bernáth, 2000<br>Csupor & Szendrei, 2012 |
| 163 | *Xanthium italicum* Moretti | sterol glycosides, flavonoids, minerals,<br>seed: fatty acid | diuretic, treat thyroid disorders | Dános Béla, 2006 |
| 164 | *Xanthium strumarium* L. | sterol glycosides, flavonoid, minerals<br>seed: fatty acid, atractyloside | diuretic, treat thyroid disorders | Dános Béla, 2006 |

**Table S2.** Classification of secondary metabolites occurred on the kurgans.

| No. | Group of secondary metabolites | Compounds / Type of compounds | References |
|---|---|---|---|
| **1** | Flavonoids | flavones, flavonols, flavanones, flavanonols, anthocyanins, chalcones | Veitch & Grayer, 2011 |
| **2** | Phenolic compounds | phenolic acids / phenolcarboxylic acids, stilbenoids, phenylethanoids, hydroxybenzyl alcohol | Kumar & Goel, 2019<br>Roupe et al., 2006<br>Abreu et al., 2011 |
| **3** | Tannins | astringent compounds, non-flavonoid polyphenols | Sharma et al., 2021 |
| **4** | Essential oil | mono- and sesquiterpenes, volatile phenolics/phenylpropanoid | Ahmad et al., 2021<br>Raut & Karuppayil, 2014 |
| **5** | Alkaloids | | Aniszewski, 2015 |
| **6** | Plant steroids | phytosterols (tetracyclic triterpenoids) | B. Gunaherath & Gunatilaka, 2014 |
| **7** | Triterpenes | Triterpenes and triterpenoids (e.g. ursolic acid, oleanolic acid) | Garg et al., 2020 |
| **8** | Polysaccharides | mucilage, starch, inulin, pectin | Bokov et al., 2020 |
| **9** | Vitamins | water-soluble and lipid-soluble vitamins | Asensi-Fabado & Munné-Bosch, 2010 |
| **10** | Other non-alkaloid nitrogen containing compounds | aminoacids<br>aminoacid derivatives, amides, amines , allantoin | Kaur et al., 2021 |
| **11** | Saponins | triterpene- saponins, steroid-saponins | Wina et al., 2005 |
| **12** | Coumarins | non-flavonoid polyphenols | Venugopala et al., 2013 |
| **13** | Organic acid | e.g. citric acid, malic acid | Huang et al., 2021 |
| **14** | Mineral nutrition | Silicon dioxide, trace elements (Fe, Mg) | Miller, 2014 |
| **15** | Fatty acids | Plant fatty acids (e.g.: oleic, linoleic, α-linolenic acids) | Lichtenstein, 2013,<br>He et al., 2020 |
| **16** | Plant proteins | | Sá et al., 2020 |
| **17** | Iridoids | bitter compounds (highly oxygenated monoterpenes) | Dinda & Dinda, 2019 |
| **18** | Lignans | di-phenolic compounds, non-flavonoid polyphenols | Xu et al., 2022 |

| | | | |
|---|---|---|---|
| **19** | Sesquiterpenes | Sesquiterpenes and sesquiterpenoids | Lorigooini et al., 2020 |
| **20** | Sesquiterpene lactones | bitter compounds | Chadwick et al., 2013 |
| **21** | Carotenoids | tetraterpenes | da Silveira Vasconcelos et al., 2020 |
| **22** | Resin | resine glycoside | Langenheim, 2003 |
| **23** | Diterpenes | Diterpenes and diterpenoids, bitter diterpenes | Eksi et al., 2020 |
| **24** | Quinones | anthraquinones, benzoquinones, naphthoquinones, naphthodianthrone | Nollet & Gutierrez-Uribe, 2018<br>Zhang et al., 2022<br>Dulo et al., 2021 |
| **25** | Steroidal glycosides | cardiac glycosides (triterpenes) | Kreis & Mller-Uri, 2010 |
| **26** | Glucosinolate | sulfur containing glycosides isothiocyanate, benzyl isothiocyanate<br>non-alkaloid nitrogen containing compounds | Abdelshafeek & El-Shamy, 2023 |
| **27** | Xanthone | tricyclic polyphenols | Remali et al., 2022<br>Vieira & Kijjoa, 2005 |
| **28** | Naphthalenes and derivatives | | Ibrahim & Mohamed, 2016 |
| **29** | Triglyceride | | Lichtenstein, 2013 |
| **30** | Phloroglucinol derivatives | non-flavonoid polyphenols | Bridi et al., 2018 |
| **31** | Cyanogenic glucosides | Cyanogenic glucosides<br>Non-alkaloid nitrogen containing compounds | Tahir et al., 2024 |
| **32** | Monosaccharides | | Stylianopoulos, 2013 |
| **33** | Phyto-cannabinoids | Terpenophenolic compounds | Brenneisen, 2007 |
